# Engineering cell morphology by CRISPR interference in *Acinetobacter baylyi* ADP1

**DOI:** 10.1101/2022.05.02.490284

**Authors:** Jin Luo, Elena Efimova, Daniel Christoph Volke, Ville Santala, Suvi Santala

## Abstract

Microbial production of intracellular compounds can be engineered by, for example, redirecting the carbon flux towards products and increasing the cell size. Potential engineering strategies include exploiting clustered regularly interspaced short palindromic repeats interference (CRISPRi)-based tools for controlling gene expression. Here, we applied CRISPRi for engineering *Acinetobacter baylyi* ADP1, a model bacterium for synthesizing intracellular storage lipids, namely wax esters. We firstly established an inducible CRISPRi system for strain ADP1, which enables tightly controlled repression of target genes. We then targeted the glyoxylate shunt to redirect carbon flow towards wax esters. Secondly, we successfully employed CRISPRi for modifying cell morphology by repressing *ftsZ*, an essential gene required for cell division, in combination with targeted knock-outs to generate significantly enlarged filamentous or spherical cells, respectively. The engineered cells sustained increased wax ester production metrics, demonstrating the potential of cell morphology engineering in the production of intracellular lipids.

## 1. Introduction

In microbial production, synthesis pathways often compete with growth for carbon and cellular resources. While static modification, such as gene deletions, can have positive effects on production by redirecting carbon fluxes, they often perturb growth-related reactions and may thus lead to unwanted trade-offs in the overall process. Strategies involving dynamic regulation of pathway fluxes have been recently developed to alleviate the constraints related to the unbalanced distribution of resources (Brockman and Prather, 2015; Hou *et al*., 2020; Wei *et al*., 2022). In our previous study, the expression of isocitrate lyase, controlled by an inducible promoter, was autonomously downregulated through growth-dependent depletion of the inducer, allowing an increased carbon flux from acetate towards lipid synthesis in *Acinetobacter baylyi* ADP1 (hereafter ADP1) (Santala *et al*., 2018).

In comparison with other targets for genetic engineering, cell morphology is often neglected but can also impact a bioprocess, particularly for the production of intracellular products (Jiang and Chen, 2016; Volke and Nikel, 2018; Wang *et al*., 2019; Kozaeva *et al*., 2021). For example, large-sized cells can provide more cellular space to accumulate intracellular products, thus allowing for a higher product content (Jiang and Chen, 2016; Zhang *et al*., 2020). In addition, cell enlargement or aggregation has been suggested to be advantageous for downstream cell recovery processes through improved filtration efficiency and the use of convenient methods, such as sedimentation and press filtration, for cell recovery from fermentation broth (Wang *et al*., 2014, 2019; Yenkie *et al*., 2017).

The variables that affect cell size have been investigated in *E. coli*, including mass doubling time, DNA replication initiation, and DNA replication-cell division cycle (Si *et al*., 2017). Controlling cell division has been demonstrated as an effective strategy to engineer cell size for bioproduction (Jiang and Chen, 2016). In recent studies, cell enlargement has been achieved by down-regulation of *ftsZ*, an essential gene required for cell division, which was shown to improve the accumulation of polymers, such as poly(lactate-co-3-hydroxybutyrate) and poly(3-hydroxybutyrate) (PHB) (Elhadi *et al*., 2016; Ding *et al*., 2020; Kozaeva *et al*., 2021). Besides, manipulation of cell shape may also affect the capacity of cells for product accumulation; for example, spherical cells have a lower surface-to-volume ratio for unit volume than rod-shaped cells. For rod-shaped species, the rod shape is typically mediated by the “rod complex”, consisting of the actin homolog MreB and other proteins, including RodA (transglycosylase) and penicillin-binding proteins (PBP) (Cho *et al*., 2016; Carballido-López, 2019).

ADP1 represents an emerging model organism for research and engineering (Metzgar, 2004; Cuff *et al*., 2012; Tumen-Velasquez *et al*., 2018; Biggs *et al*., 2020; Santala and Santala, 2021). It does not only share most of the traits that make *E. coli* a convenient model organism but also possesses unique features, such as natural competence and genetic tractability (Metzgar, 2004). In addition, the bacterium is known for its ability to accumulate high-value storage compounds-wax esters (WEs) (Martin *et al*., 2021). We have previously engineered the WE production pathway by increasing the availability of the precursor acetyl-CoA and overexpression of the WE synthesis gene (Lehtinen *et al*., 2018; Santala *et al*., 2018; Luo *et al*., 2020). WE accumulation was also facilitated by employing a strategy that partitioned the carbon sources between growth and WE synthesis (Santala *et al*., 2021). However, cell morphology engineering has not been carried out for WE production. The synthesis of WEs starts at the inner side of the cytoplasm membrane, forming membrane-bound lipid-prebodies, which are subsequently released to form matured lipid-bodies (Wältermann *et al*., 2005). It is yet to be explored how cell morphology changes affect WE body formation and whether the strategy can be applied with WEs.

A more recent engineering strategy allowing for dynamic pathway regulation involves the clustered regularly interspaced short palindromic repeats (CRISPR) system. In nature, CRISPR is widely encoded by bacteria and archaea, providing protective mechanisms against the invasion of foreign genetic elements (Makarova *et al*., 2011). One of the simplest CRISPR systems described thus far is from *Streptococcus pyogenes*, which relies on the single endonuclease Cas9. The CRISPR-Cas9 system provides RNA-guided site-specific DNA cleavage and has been repurposed as an efficient genome editing tool (Jiang *et al*., 2013). Substitution of Cas9 for its catalytically inactive mutant, referred to as dCas9, which retains DNA-binding capacity but is unable to cleave DNA, can block RNA polymerases from transcription; this synthetic system is called CRISPR interference (CRISPRi) (Qi *et al*., 2013). An inducible CRISPRi system serves as a powerful tool for repression studies of essential genes and strain optimization by co-expression of *dCas9* with a single guide RNA (sgRNA) targeting specific genes (Woolston *et al*., 2018; Guzzo *et al*., 2020). Simultaneous repression at multiple loci is enabled by the expression of multiple different sgRNAs. To this end, these sgRNAs can be transcribed in a single transcript which is subsequently processed into multiple functional sgRNAs (McCarty *et al*., 2020). For example, the endonuclease Cas6 (formerly known as Csy4) from *Pseudomonas aeruginosa* (Haurwitz *et al*., 2010) has been applied in different host strains for site-specific RNA processing (Qi *et al*., 2012; McCarty *et al*., 2019).

CRISPRi has been previously established and employed in ADP1 (Geng *et al*., 2019). The goal was to repress the expression of mobile elements, which allowed placing *dCas9* and sgRNA under the control of constitutive promoters. Here, CRISPRi was further developed in ADP1 for metabolic engineering purposes, namely for the redirection of carbon flux and engineering of cell morphology that involves essential genes as targets. To allow conditional knockdowns, inducible systems for both *dCas9* and sgRNA were established and optimized. In addition, the ribonuclease Cas6 allowing simultaneous repression of multiple loci was studied to expand the CRISPR-based toolbox for ADP1. The developed systems were then investigated in the context of cell morphology and its effects on intracellular lipid (WE) production.

## 2. Results

### 2.1. Establishment of inducible CRISPRi in *Acinetobacter baylyi* ADP1

To target growth-essential genes in ADP1, we first established an inducible CRISPRi system based on dCas9. To ensure tight control of the system, two induction systems were used to control the expression of *dCas9* and sgRNA separately (Figure 1A). An arabinose-inducible promoter was used to control the expression of sgRNA in a pBAV plasmid. The vector replicates in ADP1 with a medium copy number (∼58) (Bryksin and Matsumura, 2010). The pBAV plasmid-based arabinose induction system was characterized in our previous study (Santala, Efimova, *et al*., 2014). A cyclohexanone-inducible promoter was optimized to control the expression of *dCas9* from the genome. The genome-integrated cyclohexanone induction system was first characterized using the red fluorescent protein mScarlet with a strong ribosome binding site (RBS) BBa_0034 (iGEM Part Registry); 5 μM cyclohexanone was sufficient to fully induce the promoter. The transfer function, showing the change in promoter activity in regards to cyclohexanone concentration, is given in Figure S1, and the corresponding model for fitting the data is presented in Supplemental Note. In order to tune the expression strength, we characterized two weaker RBSs from the iGEM Part Registry, BBa_J61138 and BBa_J61105, by linking them to the mScarlet gene. These RBSs lead to ∼200 and ∼2000-fold lower fluorescence intensities than BBa_0034 upon full induction (Figure S2).

**Figure 1.**
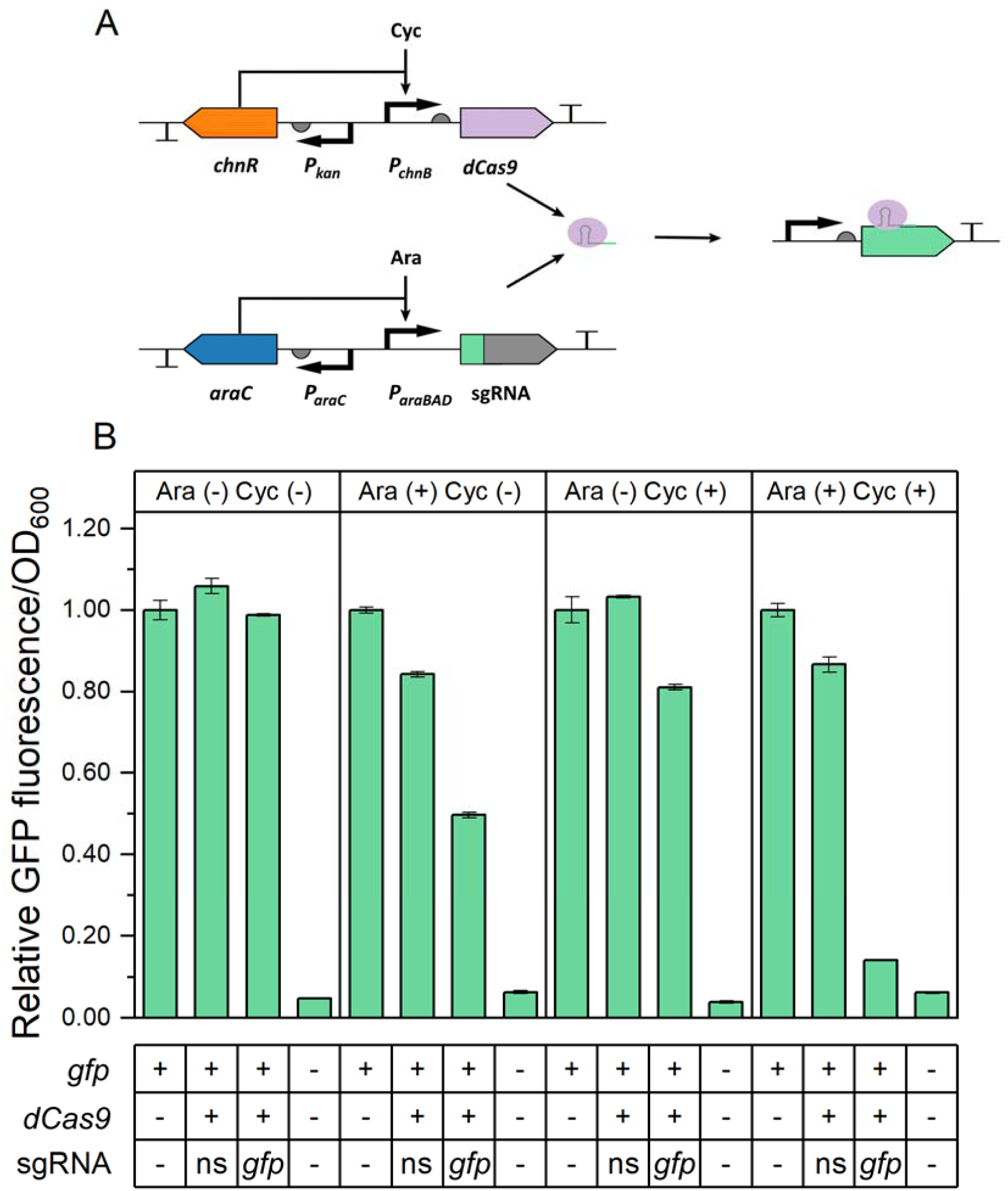
Characterization of the CRISPR interference system in *A. baylyi* ADP1. (A) Design of the CRISPRi system. The *chnR* regulator and *dCas9* are expressed from the genome, and the *araC* regulator and sgRNA are expressed from the pBAV plasmid. The *dCas9* gene and sgRNA (spacer sequence in green) are under the control of the cyclohexanone (Cyc)-inducible promoter and arabinose (Ara)-inducible promoter, respectively, while the regulators are constitutively expressed. (B) Analysis of the CRISPRi system under different induction states using GFP as the reporter. Cells were grown in LB media and cultivated at 30 L. For each condition, the data at 26 h is shown as the fluorescence/OD_600_ relative to the values from the GFP reporter strain without sgRNA and dCas9. The time-dependent change of the fluorescence signal is presented in Supplemental Materials (Figure S4). As controls, wild-type ADP1 and the GPF reporter strain containing *dCas9* and a non-specific (ns) sgRNA are used. Data represent average values ± standard deviations of two independent biological experiments.

To evaluate the efficiency of the system, we tested the repression in an msfGFP (hereafter GFP) reporter strain (ASA514). The strong constitutive promoter *P*_*14g*_ was used to drive *gfp* expression from the genome (Zobel *et al*., 2015). In an initial experiment, the strong RBS BBa_0034 was used to drive the expression of *dCas9*. This led to a greatly reduced fluorescence signal, upon full induction of the expressions of the *gfp*-targeting sgRNA (targeting base 41-60 of the coding region) and *dCas9* with 1% (w/v) arabinose and 5 μM cyclohexanone. However, the growth rate decreased notably. This is likely due to the toxicity of strongly expressed *dCas9*, stemming from its non-specific binding to NGG sequences in the genome (Zhang and Voigt, 2018). Although the levels of dCas9 can be reduced by lowering the inducer concentration (Figure S1), the low transcription and high translation rate would lead to high expression noise and in turn to phenotypic variation across the cell population and potentially raise uncertainties (Mundt *et al*., 2018). Additionally, the strong RBS leads to higher basal expression and therefore to repression without induction. Thus, the strong RBS preceding *dCas9* was replaced by BBa_J61138 to reduce the dCas9 dose when fully induced. Upon full induction, the strain with the weak RBS had a 3.3% reduction in growth rate, while the strain with the strong RBS exhibited a 32% reduction (Figure S3). The lower *dCas9* expression level still enabled effective gene repression; by inducing the expression of *dCas9* and the *gfp*-targeting sgRNA, the fluorescence of the strain decreased 86% compared to the reporter strain without the CRISPRi machinery (Figure 1B).

Effective repression was maintained throughout the whole cultivation (Figure S4). Induction with either inducer led to different repression folds, emphasizing the importance of using inducible promoters for *dCas9* and sgRNA.

### 2.2. Multiplex CRISPRi mediated by the ribonuclease Cas6

Motivated by the effectiveness of the single-guided CRISPRi, we set out to expand the system for multiplexed repression, by using several sgRNA simultaneously. To this end, we employed Cas6 for processing a guide RNA array. Cas6 (formerly known as Csy4) is a ribonuclease of the CRISPR system in *P. aeruginosa* that processes primary CRISPR transcripts (Haurwitz *et al*., 2010). Two sgRNAs targeting *gfp* and *mScarlet* were combined in the guide RNA array (Figure 2A), which targeted base 41-60 and base 142-164 of the corresponding coding sequence. The 20-bp Cas6 recognition sites were placed upstream of each sgRNA and downstream of the second sgRNA. On the same pBAV plasmid, the *cas6* gene was expressed from the constitutive promoter BBa_J23150 (iGEM part). The two-sgRNA array was tested in a double reporter strain ASA518 containing *dCas9* under the transcriptional control of the cyclohexanone-inducible promoter. As shown in Figure 2B and Figure 2C, the msfGPF and mScarlet fluorescence was repressed by 69% and 61% respectively upon induction with 1% (w/v) arabinose and 5 μM cyclohexanone, indicating that the Cas6-mediated multiplex CRISPRi was well-functioning. Interestingly, in the control experiment, the strain lacking *cas6* also showed reduced GFP fluorescence upon induction but not for mScarlet (Figures 2B and (C), indicating that the unprocessed gRNA array may still be effective in guiding the repression of the gene targeted by the first sgRNA. It seems like the function of the first sgRNA in the unprocessed array, which targeted *gfp*, is neither affected by the 32 bases preceding the spacer (+1 site till spacer) nor by the succeeding second sgRNA (Figure S5). To further validate this, we tested the plasmid containing the guide RNA array but lacking *cas6* in the GFP single reporter strain; a clear reduction of fluorescence signal was observed upon induction (Figure S6).

**Figure 2.**
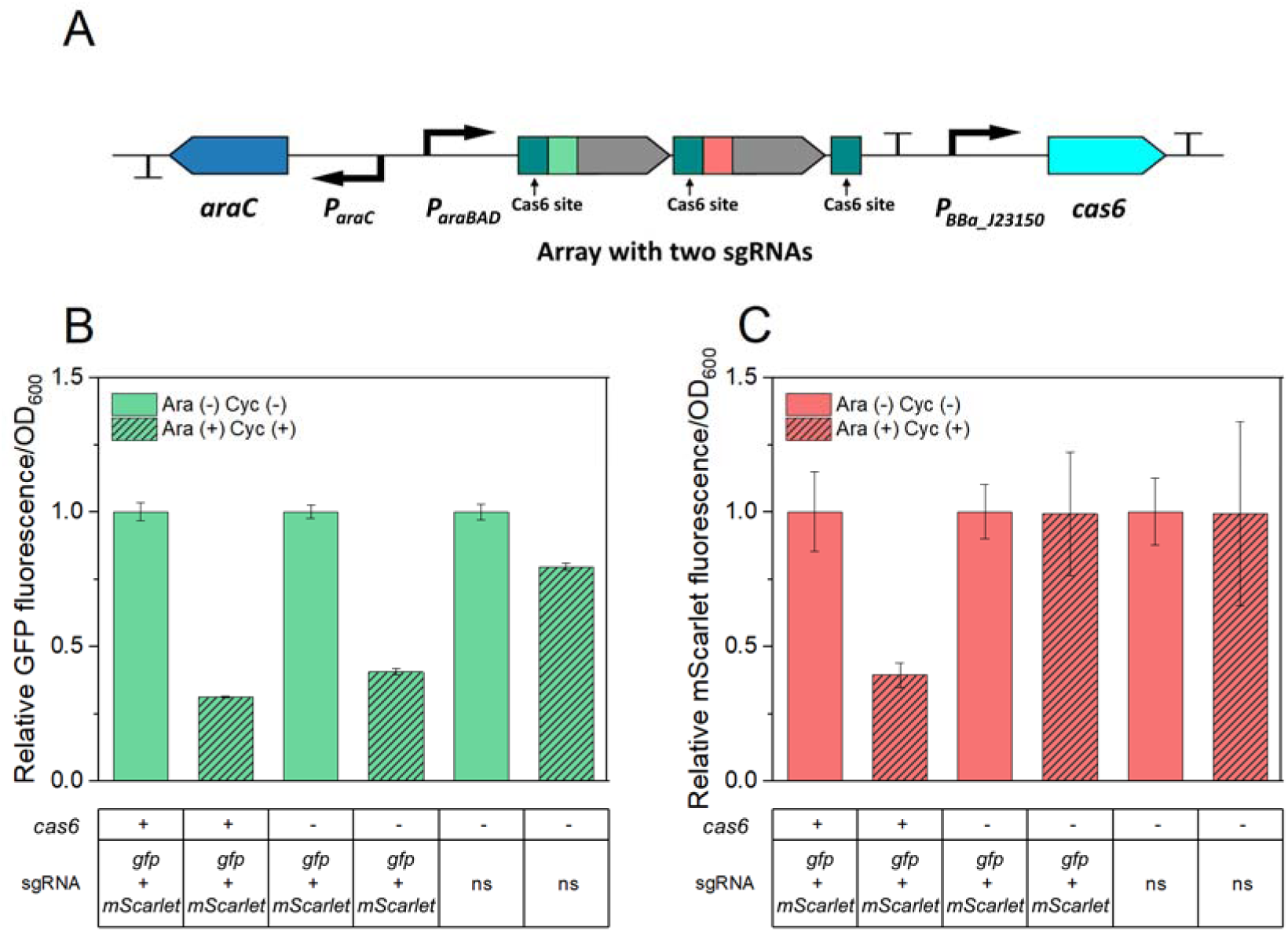
Characterization of the Cas6-mediated multiplex CRISPRi. (A) Design of the two-sgRNA array expressed in the pBAV plasmid. A Cas6 recognition site is placed before each sgRNA spacer sequence (in green and red). The third Cas6 recognition site is placed behind the second sgRNA. The expression of *cas6* is controlled by a constitutive promoter (iGEM part: BBa_J23150). Simultaneous repression of *gfp* (B) and *mScarlet* (C) by expression of the two-sgRNA array in the double reporter strain containing *dCas9*. Cells were grown in buffered LB containing 0.4% (w/v) glucose and cultivated at 30 L. Samples were taken for fluorescence measurement after 20 h of cultivation. The fluorescence values for each strain were normalized to the values measured for this strain without being induced by cyclohexanone (Cyc) and arabinose (Ara). Two *Cas6*-negative strains were used as the controls, which carried the two-sgRNA array and a non-specific (ns) sgRNA, respectively. Data represent average values ± standard deviations of two independent biological experiments.

### 2.3. Applying the CRISPRi toolset for metabolic engineering

#### 2.3.1. Modulating fluxes through the glyoxylate shunt by *aceA* repression

We next sought to demonstrate the use of the CRISPRi system for pathway modulation. The glyoxylate shunt replenishes the tricarboxylic acid (TCA) cycle intermediates from acetyl-CoA, bypassing the two oxidative steps that release two molecules of carbon dioxide (Figure 3A). The shunt is essential for biomass synthesis when acetate is used as the sole carbon source. Isocitrate lyase, AceA, catalyzes the first step of the glyoxylate shunt. Previously, it has been shown that in ADP1 gradual downregulation of *aceA* led to decreased biomass formation from acetate and redirection of the carbon flux towards fatty acid synthesis, thus enhancing WE accumulation (Santala *et al*., 2018). Here, we tested the repression of *aceA* by CRISPRi. We grew cells in MA/9 minimal media supplemented with casamino acids (0.2% W/V) and 40 mM acetate at 30 L. For induction, 1% (w/v) arabinose and 5 μM cyclohexanone were supplemented. Wild-type ADP1 showed slightly faster growth and acetate consumption than the strains containing the CRISPRi system (Figure 3B). In the absence of inducers, there was no difference in growth and acetate consumption between the two CRISPRi strains targeting either *aceA* or GFP gene; upon induction, the strain targeting *aceA* showed slower growth and acetate consumption and a lower final OD (Figure 3B), indicating effective repression of the glyoxylate shunt for acetate metabolism. In *aceA*-repressed cells, WEs were detected after the 20 h of cultivation (Figure 3C), while WEs were not observed in the control strains (without *aceA* repression).

**Figure 3.**
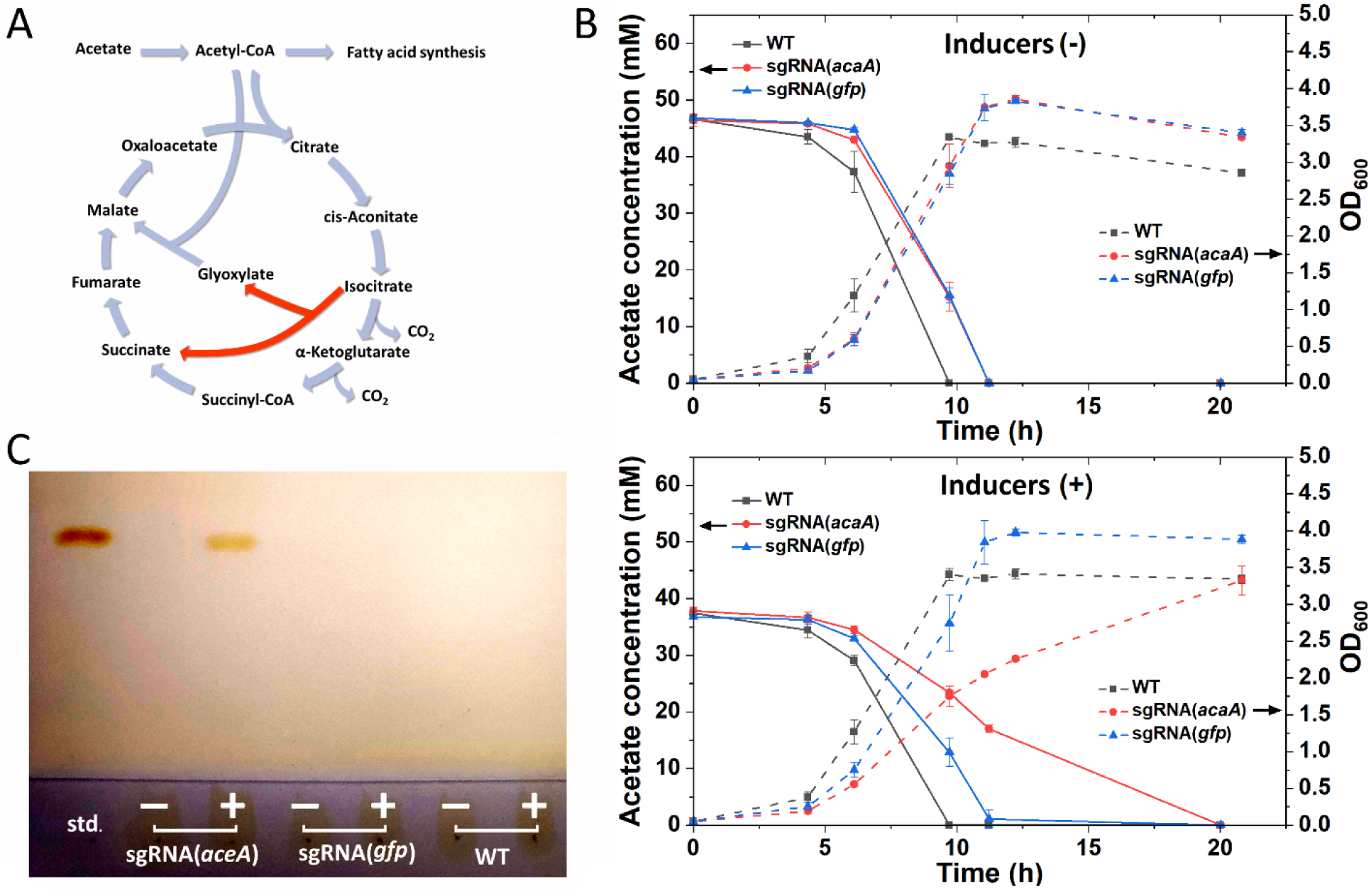
Repression of *aceA* by CRISPRi. (A) Schematic of the TCA cycle and glyoxylate shunt. The glyoxylate shunt replenishes the TCA cycle intermediates using two molecules of acetate as substrates and is thus essential for cells grown on acetate. The enzyme AceA catalyzes the first reaction of the glyoxylate shunt (red). (B) The CRISPRi strain carrying the *aceA*-targeting sgRNA showed lower growth and acetate consumption rates upon induction than the control strain containing the GFP gene-targeting sgRNA. Wild-type (WT) ADP1 was used as a control. All the strains were cultivated in MA/9 media supplemented with 40 mM acetate and 0.2% (w/v) casamino acids at 30 L. Data represent average values ± standard deviations of two independent biological experiments. (C) TLC was used to analyze WEs from the biomass at the endpoint (20 h) for each strain with (+) and without (-) inducers. Jojoba oil was used as the standard (std.).

#### 2.3.2. Manipulating cell morphology by *ftsZ* repression

Cells with large size and/or atypical morphology can have benefits for intracellular product accumulation and downstream processing, such as higher storage capacity and easier cell harvesting (Jiang and Chen, 2016; Wang *et al*., 2019). Cell division is one of the processes that can be manipulated to control cell size: a shortened division period decreases cell size, and a prolonged division period leads to cell elongation. To realize cell enlargement in APD1, we applied the CRISPRi system to repress *ftsZ*, one of the essential genes for cell division (Figure 4A), and thereby increase cell size. We also sought to explore the effect of *ftsZ* repression in the spherical mutants where the rod system was inactivated (Figure 4A).

**Figure 4.**
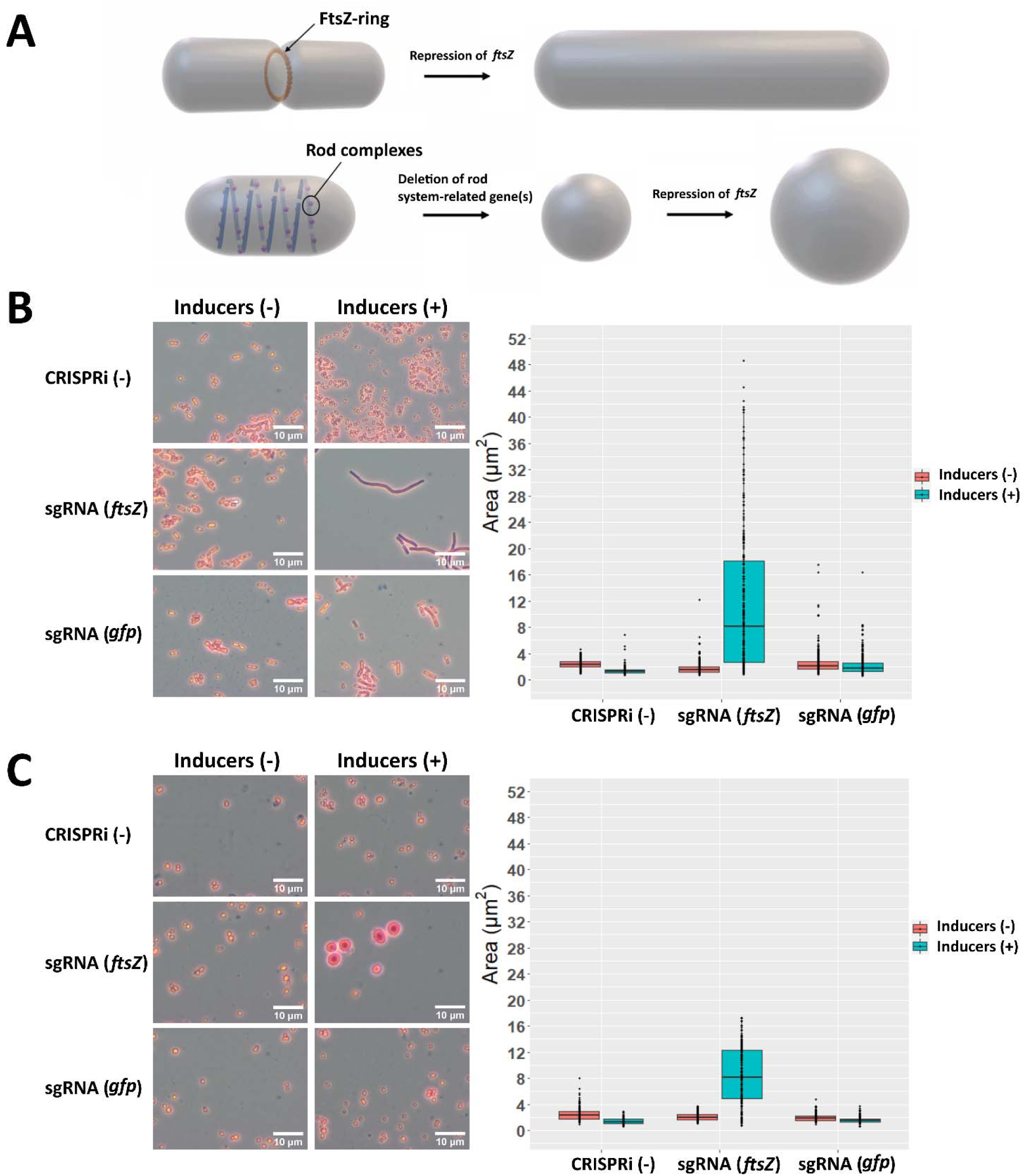
Morphological changes and increase of cell size by *ftsZ* repression. (A) Strategies for engineering cell size and shape. The FtsZ-ring is essential for cell division. It acts as the scaffold for recruiting other division proteins and provides contractile force to divide the daughter cells. One model indicates that the MreB polymerizes into long filaments and binds to other peptidoglycan synthesis proteins, such as PBP2 (PbpA) and RodA, to form “rod complexes”. Deletion of the rod-system-related gene(s) leads to spherical cells. Repression of *ftsZ* in the normal rod-shaped strain led to cell filamentation (B), while *ftsZ* repression in the *pbpA* mutant led to enlarged spherical cells (C). Cells were cultivated in LB media at 30 L. The strains without the CRISPRi machinery and the strains carrying the *gfp*-targeting sgRNA served as controls. Cells were stained with crystal violet before being viewed under the microscope. Two independent biological experiments were performed, from which similar cell morphology changes were observed under the microscope. Photos were shown, and cell size quantification was done for one of the experiments. For each sample, 200 or 300 cells were picked randomly for size analysis, which is shown as the cross-section areas of cells on the images.

We first validated the strategy by treating a rod-shaped strain and two spherical strains (deletions of *mreB* or *pbpA*) with piperacillin. Piperacillin is a penicillin beta-lactam antibiotic that inhibits PBP3, a central component of cell divisome (Kocaoglu and Carlson, 2015). As expected, after piperacillin treatment, cell filamentation was observed for the rod-shaped strain, and large spherical morphology was observed for the *mreB* and *pbpA* mutants (Figure S7). We then introduced the CRISPRi system targeting *ftsZ* into wild-type ADP1 and a *pbpA* deletion mutant. The strains carrying CRISPRi targeting *gfp* and the strains without the CRISPRi system served as control strains. We grew cells in LB media and cultivated them at 30 L. Inducers (1% (w/v) arabinose and 5 μM cyclohexanone) were added when cells were in the early exponential phase. Eighteen hours after induction, cells were investigated through the microscope, and the cell size was determined through the cross-section of single cells on the image. The induction led to an increase in cell size for both, the rod-shaped and spherical strains targeting *ftsZ*. For both strains, the median of cell cross-section for randomly selected cell populations under the microscope increased by more than four-fold (Figures 4B and (C). Noteworthy, a larger variance in cell size was observed for the population of the rod-shaped strain. Interestingly, in the presence of inducers, the rod-shaped strain continued to elongate as long as nutrients were not limited, while the spherical strain stopped growth at a certain size even with an excess of nutrients. Thus, larger cell size can be obtained in rod-shaped cells than in spherical cells by *ftsZ* repression, and the increment of cell size can be controlled through the induction at different growth stages.

Cell size can influence sedimentation (Wang *et al*., 2019). A larger cell settles faster, given that the density of the single cell is not reduced and is higher than the surrounding fluid. On this background, we then compared the sedimentation of the cells with and without manipulation of cell enlargement. Both, in the rod-shaped and the spherical strains, cell enlargement led to faster sedimentation (Figure 5B).

**Figure 5.**
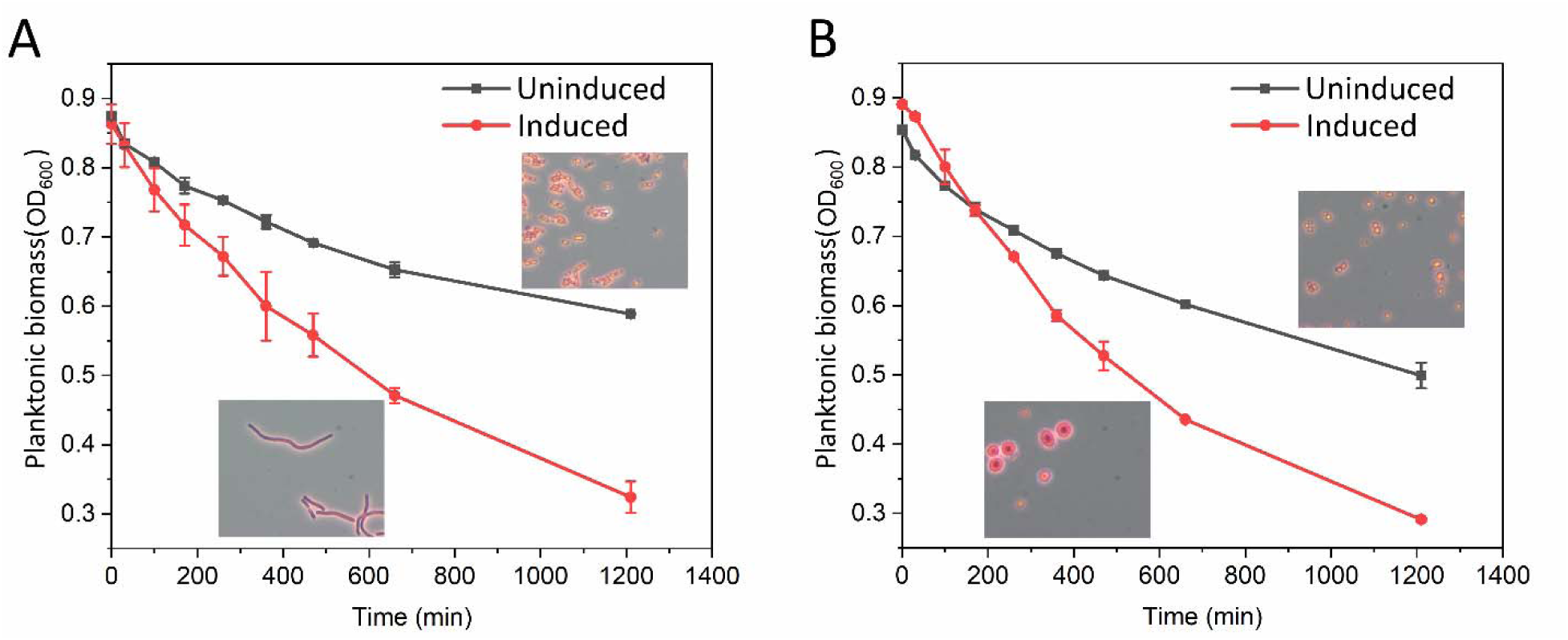
Sedimentation of enlarged cells. Sedimentation in both strains, the rod-shaped strain (A) and the *pbpA* mutant (B), was analyzed with and without induction of CRISPRi for repression of *ftsZ*. After cultivation in LB media, cells were washed and resuspended in PBS in test tubes. Microscope pictures are taken from (Figure 4. Two independent sedimentation experiments were performed with the cells from the same culture. Data represent average values ± standard deviations of the two independent sedimentation experiments.

### 2.4. Cell enlargement in different cultivation conditions for WE production

#### 2.4.1 Cultivation by co-feeding of glucose and acetate or using only glucose

We next aimed to study the production of WEs in enlarged cells. Although wild-type ADP1 naturally produces wax esters as storage compounds, the production metrics are low. Previously, co-feeding of a glycolytic carbon source, gluconate, and acetate was performed with ADP1 Δ*aceA* overexpressing *acr1* (Figure 5A); the strategy enabled partition of carbon source metabolism such that the glycolytic substrate was used for cell growth while acetate could be directed for WE synthesis, allowing enhanced WE accumulation (Santala *et al*., 2021). It has also been shown that *aceA* deletion and *acr1* overexpression could lead to improved WE production with high glucose supplementation in the media (Luo *et al*., 2020). We applied the same engineering strategy in both the rod-shaped strain and the spherical *pbpA* mutant. The CRISPRi system targeting *ftsZ* was further introduced into the modified strains. The two resulting strains were designated as ASA531 (Δ*aceA* + *acr1* overexpression + CRISPRi targeting *ftsZ*) and ASA532 (Δ*pbpA* + Δ*aceA* + *acr1* overexpression + CRISPRi targeting *ftsZ*). However, it was found that ASA532 had an inconsistent and slower growth than ASA531 when glucose or acetate were supplied in high concentrations (data not shown), thus we focused on ASA531 in the succeeding experiments. MSM was used in all the experiments for WE production.

We first cultivated ASA531 using glucose and acetate as carbon sources; it is expected that cell growth is fueled by glucose, while WE synthesis is fed by acetate. To ensure a high WE content, we kept a high acetate to glucose ratio; acetate and glucose were added in the beginning with a concentration of 60 mM and 5 mM, respectively, and were supplied in the same concentrations at 10 h and 21 h to avoid depletion of carbon and other components necessary for cellular functions. Three cultures were grown in parallel, in two of which 1% (w/v) arabinose and 50 μM cyclohexanone were added when the OD_600_ of the cells reached ∼0.5 (early induction) or ∼1.5 (late induction). After 34 h of cultivation, both cultures with induction contained larger cells than the control; and cells with early induction were the most elongated (Figure 5B), suggesting that the increase in size can be controlled by the time point of induction. The cellular WE content and titer were found to be approximately 60% higher in the cells with early induction compared to the cells without induction, but the overall WE content for all the cultures was low, less than 0.06 g/g cell dry weight (CDW) (Table 1). Looking at the fermentation profiles, the biomass already started to decline for all the cultures after the second carbon source supplementation (∼21 h) (Figure S8); meanwhile, glucose consumption almost stopped, and acetate consumption was reduced after 21 h (Figure S8). At the end of the cultivation, all cells consumed 9 mM glucose in total. For the cells without induction, with early induction, and with late induction, the total acetate consumption was 115 mM, 139 mM, and 133 mM, respectively (Figure S8).

**Table 1.**
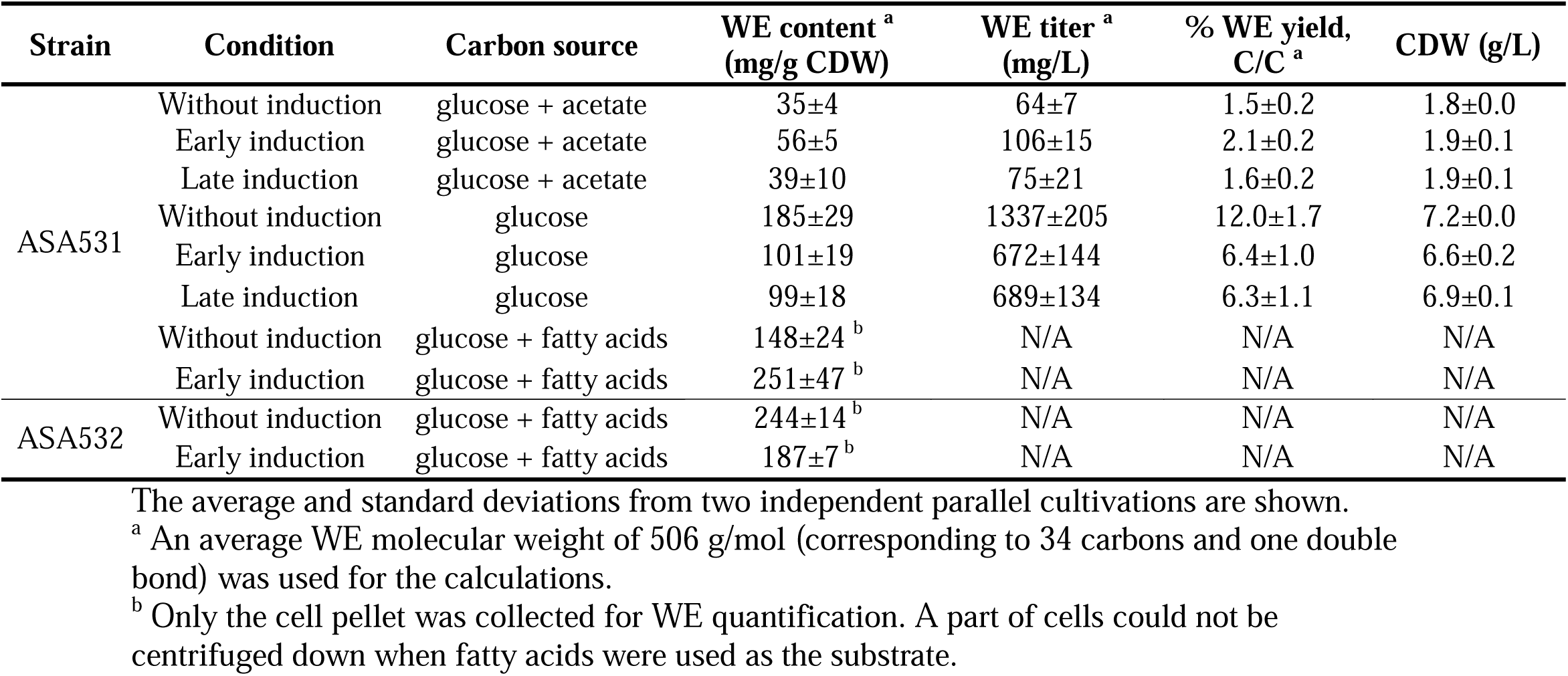
WE production by ASA531 and ASA532.

Therefore, we next cultivated cells in high glucose concentration (200 mM) as this enables cultures to reach higher final biomass concentrations. Induction was performed at higher ODs (OD of ∼1.5 for early induction and ∼4 for late induction). After 48 h of cultivation, 124 mM, 118 mM, and 123 mM glucose were consumed by the cells without induction, with early induction, and with late induction, respectively. In these conditions, the overall WE production was improved for both cultures with and without induction, although the cells with induction produced less WEs (∼0.100 g/g CDW) compared to the cells without induction (0.185 g/g CDW) (Table 1). However, microscopic analyses (Figure 5B) revealed successful increase of cell size for the cells with induction, thus indicating a higher storage capacity and a potential to further enhance WE accumulation in the enlarged cells.

#### 2.4.2 Cultivation by co-feeding glucose and fatty acids

*De novo* synthesis of WEs from glucose or acetate involves multiple steps. In addition, the fatty acid synthesis pathway is under tight regulation (Janßen and Steinbüchel, 2014). All these pose a hindrance to WE accumulation. We hypothesized that by using fatty acids (FAs) as the substrate for WE synthesis (Figure 5A), it would be possible to bypass the long synthesis pathway and regulation, and by that circumventing potential inherent limitations for WE synthesis. To this end, we used FAs as the substrate to investigate whether the enlarged cells were capable of storing larger amounts of WEs. In addition, FAs are attractive substrates for WE synthesis as they can be obtained from the hydrolysis of waste cooking oils. Both, ASA531 (Δ*aceA* + *acr1* overexpression + CRISPRi targeting *ftsZ*) and ASA532 (Δ*pbpA* + Δ*aceA* + *acr1* overexpression + CRISPRi targeting *ftsZ*), were cultivated in media containing 10 mM glucose and ∼14 g/L FAs (mainly oleic acid: 65%-88%). At 12 h, 10 mM glucose was supplemented, and inducers were added for cell enlargement. After 34 h of cultivation, the enlarged cells were found to accumulate a large amount of lipid droplet, observed by microscopic analysis (Figure 5B). The identity of the lipid droplets was validated by Nile red staining (Figure 5B). Interestingly, for both cultures with and without induction, a part of the cells could not be sedimented by centrifugation and was suspended in or floating on top of the media (Figure S9). This could result from the use of fatty acids as the substrate, which may reduce the density of single cells (or cell-matrix). The density can also be reduced due to WE accumulation. Eventually, only the part of the cells that were collected by centrifugation was subjected to WE quantification. According to WE quantification, the enlarged cells of ASA531 accumulated WEs up to 0.251 g/g CDW, which was higher than the content of the cells without induction (0.148 g/g CDW; Table 1). However, the enlarged ASA532 had lower WE content (0.187 g/g CDW) than the ASA532 cells without induction (0.244 g/g CDW; Table 1). The WE content might be slightly underestimated as we used the previously determined the average molecular weight of WEs of 506 g/mol, corresponding to 34 carbons (Lehtinen *et al*., 2018). However, the supplementation of the fatty acid substrate, mostly consisting of oleic acid (36 carbons), may increase the average WE molecular weight.

**Figure 5.**
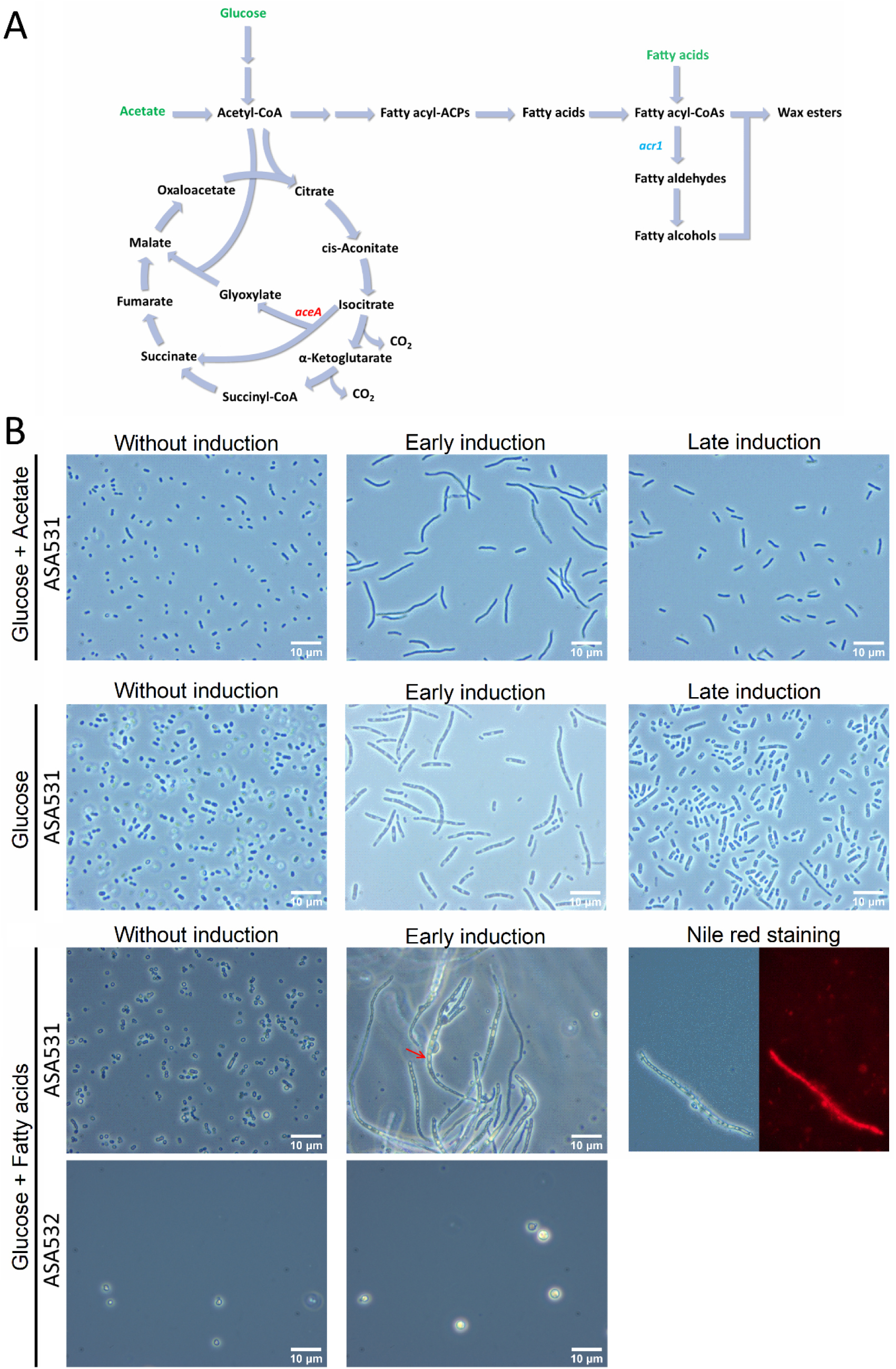
WE production in enlarged cells. (A) Schematic of WE synthesis pathway in ADP1. The substrates used in WE production are highlighted in green. In the WE production strains used in this study, *aceA*, which encodes isocitrate lyase, was deleted, and *acr1*, which encodes fatty acyl-CoA reductase, was overexpressed. Double arrows indicate that the pathway involves multiple reactions. (B) The effect of *ftsZ* repression on cell morphology during the cultivations of ASA531 (*aceA* deletion + *acr1* overexpression + CRISPRi targeting *ftsZ*) and ASA532 (Δ*pbpA* + *aceA* deletion + *acr1* overexpression + CRISPRi targeting *ftsZ*) with different carbon sources. Lipid droplets (red arrow) were visible in the enlarged cells cultivated with fatty acids under the microscope. Nile red was used to stain the lipid body.

## 3. Discussion

The development of cell factories that meet economic requirements for industrial production needs significant research efforts. For intracellular products, it can be even more challenging due to the high cost of product recovery. The cost of the downstream processes, which can contribute to more than 70% of the total production cost, can be significantly reduced if high biomass titer and product content are achieved, and thus less effort is needed for the biomass and product collection (Yenkie *et al*., 2017). However, the accumulation of intracellular products can be limited due to the small size of bacterial cells. Larger cells provide more space for product accumulation and can thus be more efficient for production (Jiang and Chen, 2016; Wang *et al*., 2019). In addition, cell size can have a great influence on cell harvesting. For example, large-sized cells may be more effectively separated by filtration or enable a faster sedimentation rate (Yenkie *et al*., 2017; Wang *et al*., 2019).

Here, inducible CRISPRi offers a convenient way to modulate gene expression for cell size control, as this modulation often reduce the fitness of the cells severely and therefore a permanent change is calamitous. Furthermore, the system is well suited for a broad range of engineering purposes and functional studies for essential genes. Previously, CRISPRi was used in *A. baylyi* ADP1 to repress the activity of IS elements, enabling a reduction of mutation rate (Geng *et al*., 2019). However, both *dCas9* and the sgRNAs were constitutively expressed, which limits its application to non-essential genes. To suppress cell division, we first established an inducible CRISPRi system. The sgRNA was expressed in a medium-copy plasmid pBAV (Bryksin and Matsumura, 2010), while *dCas9* was integrated into the genome to reduce plasmid size and alleviate its potential toxicity caused by high dosage (Zhang and Voigt, 2018). The spacer sequence in the sgRNA can be easily replaced by PCR-based methods. In the previously described inducible CRISPRi systems, an inducible promoter was usually used to control the expression of dCas9 while sgRNA was expressed constitutively (Qi *et al*., 2013; Li *et al*., 2016; Peters *et al*., 2016; Woolston *et al*., 2018; Guzzo *et al*., 2020). However, such systems can lead to leaky repression (Li *et al*., 2016; Peters *et al*., 2016). Here, to minimize the leakage, we used two inducible promoters, namely the arabinose-inducible promoter *P*_*araBAD*_ and the cyclohexanone-inducible promoter *P*_*chnB*_, to control the expression of the sgRNA and *dCas9* separately. The arabinose-inducible promoter is widely used for the expression of heterologous genes. In ADP1, arabinose can be unselectively oxidized by glucose dehydrogenase in the absence of glucose (Kannisto *et al*., 2015; Santala *et al*., 2018), which may reduce the effect of induction. This feature has been exploited to create an autonomously regulated switch for gene expression regulation (Santala *et al*., 2018) and could also be coupled with CRISPRi to make it even more dynamic. On the other hand, the oxidation of arabinose can be readily relieved by supplementing cells with glucose. The cyclohexanone-inducible promoter is originated from a cluster of genes responsible for cyclohexanol degradation in *A. johnsonii*, regulated by its cognate activator ChnR (Steigedal and Valla, 2008).

Studying essential genes by CRISPRi is highly sensitive to perturbations; leakage of the repression system, for example, can severely impair cell growth. We showed that the employment of the two induction systems enables a tightly controlled CRISRPi system. The established CRISPRi exhibits minimal leaky repression, as indicated by the unchanged phenotypes in the absence of inducers. In addition, the burden of CRISPRi was significantly reduced by optimizing the dCas9 dosage. The minimized burden is important for gene repression study, especially when exploring how the repression of a specific gene affects cell morphology, which can be easily influenced by cell physiological conditions (Si *et al*., 2017).

The ability to simultaneously target multiple loci will expand the application of CRISPRi. Multiplex CRISPRi enables systematic genetic interaction studies (Guzzo *et al*., 2020). To enhance metabolic flux towards desired products by metabolic engineering, modulation of the expression of multiple genes is often required. For example, simultaneous repression of four competing pathway genes increased the yield and productivity of n-butanol in *E. coli* (Kim *et al*., 2017). In addition, the expression of multiple sgRNAs targeting the same gene can improve the repression efficiency for this gene (McCarty *et al*., 2019). Thus, we were keen to explore multiplex CRISPRi in ADP1. Targeting multiple loci requires the expression of multiple sgRNAs. This can be accomplished by the expression of each sgRNA using an individual promoter. Alternatively, the expression platform can be composed of a single promoter and a guide RNA array, which is processed into multiple functional sgRNAs. The former strategy is straightforward and has been used in many studies (Kim *et al*., 2016; Geng *et al*., 2019; Guzzo *et al*., 2020). However, repeated use of the same promoter may trigger genetic instability due to homologous recombination. Thus, we chose to use one promoter to express a single guide RNA array encoding multiple sgRNAs. Processing of the single transcript into functional sgRNAs was achieved by co-expression of the ribonuclease Cas6 (Haurwitz *et al*., 2010), which cleaves RNA on specific sites inserted between sgRNA sequences. Cas6 is part of the native CRISPR system in *P. aeruginosa*, responsible for processing CRISPR transcripts. Besides, Cas6 functions in various other organisms, including *E. coli, Bacillus subtilis*, and *Saccharomyces cerevisiae* (Qi *et al*., 2012). Other strategies for processing guide RNA arrays have been summarized in (McCarty *et al*., 2020), such as the employment of a CRISPR effector possessing RNase activity, e.g., Cas12a. Here, we established a multiplex CRISPRi system in ADP1 for the first time. The system was successfully used to simultaneously repress *gfp* and *mScarlet*. Besides, it was found that the guide RNA array was still effective in guiding the repression of the targeted gene by the first sgRNA in the array even in the absence of Cas6. The observation indicated that the function of the first sgRNA sequence in the unprocessed transcript was not affected by the 32 bp upstream sequence and the second sgRNA sequence. A study in HEK293FT cells showed that extension of sgRNA guiding sequences beyond the 20 bp did not influence targeting specificity, which was due to the sgRNA being processed to contain the 20 bp guiding sequence (Ran *et al*., 2013). However, it remains unclear how the sequence of the second sgRNA affects the function of the first sgRNA in the unprocessed array.

The utility of the CRISPRi to modulate cell metabolism in ADP1 was first evaluated by repressing the expression of *aceA*, an isocitrate lyase catalyzing the first reaction of the glyoxylate shunt. The significance of the glyoxylate shunt downregulation in metabolic engineering has been shown in our previous study, where the carbon flux could be redirected from biomass formation to wax ester synthesis when acetate was used as the substrate (Santala *et al*., 2018). We observed that *aceA* repression specifically decreased acetate consumption and cell growth rates when cells were grown in the media containing casamino acids and acetate, indicating a lowered carbon flux from acetate to biomass and direction of carbon to WEs. The results suggest that the CRISPRi system could be used as an efficient tool for pathway optimization and conditional regulation of gene expression in metabolic engineering.

For increasing the cell size, we conditionally suppressed cell division by repressing *ftsZ* expression using inducible CRISPRi. FtsZ is critical for cell division and has been used as an effective target for cell size engineering in *E. coli* (Elhadi *et al*., 2016; Jiang and Chen, 2016; Ding *et al*., 2020). Suppression of cell division can also be achieved by overexpression of the genes coding negative regulatory proteins such as SulA and MinC (Wang *et al*., 2014; Ding *et al*., 2020), both inhibiting FtsZ assembly. Here, *ftsZ* repression resulted in cell elongation for the rod-shaped strain and enlarged spherical cells for the spherical strain (with *pbpA* deletion). Notably, the induction was effective throughout the whole population, leading to an increase in cell size for the majority of the cell population. Compared to repression in the spherical strain, repression in the rod-shaped strain is more advantageous; if induction began at lower cell densities, the rod-shaped strain could reach a larger cell size and still maintain a final biomass concentration close to the culture without induction, as the cells could continue elongating. In contrast, the spherical strain would stop growing after the cells reached a certain size, and consequently, the final biomass concentration was lower if induction was carried out in the early stage. Thus, for the spherical strain, proper timing for induction is important to achieve optimal cell size and biomass quantity at the same time.

A larger cell size can be advantageous for downstream separation processes; for example, it improves the efficiency of both filtration and gravity sedimentation (Yenkie *et al*., 2017; Wang *et al*., 2019). Elongated *E. coli* cells, engineered to accumulated polyhydroxyalkanoates (PHAs) are able to settle after 20 min (Wang *et al*., 2014). In the current study, both the elongated rod-shaped cells and the enlarged spherical cells settled faster than the normal-sized cells because the settling velocity is proportional to the square of particle radius according to Stokes’ law. It should be noted, however, that the difference in densities between the microorganisms and the surrounding fluid is also an important factor affecting the settling velocity, and the density of microorganisms can be affected by the accumulated intracellular products. Accumulation of products that are denser than the cells, such as PHB, glycogen, and sulfur, will increase the density of microorganisms (Mas *et al*., 1985), thus increasing the settling velocity. In contrast, the accumulation of lighter products, such as WEs in the current study, can decrease the density of cells, and sedimentation may not benefit from large-sized cells.

WEs are natural storage compounds accumulated by ADP1and of high economic value. Their synthesis is initiated at the cytoplasm membrane, followed by the formation of matured lipid bodies. WE or other neutral lipid synthesis has not been previously investigated in cells with altered morphology. In previous studies, WEs have been proposed to be even toxic in some hosts (Kaiser *et al*., 2013), potentially due to the lack of proper organization of the lipid bodies in the cells. Therefore, it was unclear whether the cells with altered morphology could sustain WE production. Interestingly, all the engineered cells could produce WEs intracellularly, including strains with the upregulated WE synthesis pathway. In the cultivation with co-fed glucose and acetate, the enlarged cells showed slightly higher WE content, though both the normal-sized cells and the elongated cells had low WE contents. This might have to do with cell lysis, as it was observed that the OD of the cultures had already declined to a fair extent when cells were harvested for WE quantification. While acetate was present in excess throughout the cultivation, glucose was almost depleted at 21 h. Rather than the morphology changes, the depletion of glucose might be the reason for cell lysis because the cellular metabolism is not sustained, as acetate alone does not support cell growth due to *aceA* deletion. Although glucose was supplemented again at 21 h, it was not consumed by the cells. It is thus important to employ a constant feeding strategy to avoid nutrient depletion, as has been done previously (Santala *et al*., 2021). We then cultivated the cells with high glucose concentration; although the WE content of the elongated cells was increased to ∼10% (W/W), it was evident that WE accumulation in the enlarged cells could still be increased.

To further investigate the capacity of the enlarged cells to accumulate larger amounts of WEs, we used a shortcut synthesis pathway to WEs by directly providing cells with FAs. FAs can be easily obtained by hydrolysis of waste cooking oils, which could offer a feasible feedstock for the biological production of high-value chemicals. The use of FAs enhanced WE accumulation in the elongated cells, as indicated by the formation of a large amount of lipid body that was visible under the microscope and confirmed by Nile red staining. Furthermore, WE accumulation displayed heterogeneity between elongated cells; while some cells bulged due to lipid accumulation, others still held potential storage capacity. Despite the drastic phenotypic change, the elongated cells sustained efficient WE synthesis, highlighting the advantage of large cells for accumulating intracellular products; the elongated cells (ASA531 with induction) were found to accumulate ∼70% more WEs than the normal-sized cells, despite that only part of the cells could be centrifuged and analyzed. Thus, a suitable cell collection method is still required to properly evaluate the WE content for the whole cell population. Nevertheless, it was concluded that elongated cells work better than enlarged spherical cells with respect to WE production, though enlarged spherical cells might also be advantageous in other applications. Although a decreased surface-area-to-volume ratio could allow a higher product content, a too low surface-area-to-volume ratio, which can be the case for enlarged spherical cells, may adversely affect product formation due to insufficient nutrient acquisition and transport. Overall, the current study demonstrated cell morphology engineering as a potential strategy for improving WE production.

## 4. Conclusion

In this study, an inducible and tightly controlled CRISPRi system was established in *A. baylyi* ADP1, allowing the investigation and repression of growth-essential genes for desired phenotype changes. To expand the application of the CRISPRi system, multiplex gene repression was explored by using a multiple sgRNA expression strategy based on the ribonuclease Cas6. The established CRISPRi system was further used for pathway modulation and cell morphology engineering. The cell size of *A. baylyi* ADP1 was increased by repressing *ftsZ*, an essential gene for cell division. Despite the cell division being severely perturbed by CRISPRi, the engineered cells were able to sustain high WE production and accumulated up to 25% WEs of cell dry weight. Furthermore, the microscopic analyses indicated that even higher storage capacity could be harnessed for WE accumulation. Our results demonstrate the benefit of the inducible CRISPRi system for a variety of applications, including genome-scale gene functional study and metabolic engineering in ADP1, and warrant further studies on large-sized cells for the production of the high-value intracellular product, wax esters.

## 5. Experimental procedures

### 5.1 Strains and media

*E. coli* XL1-Blue (Stratagene, USA) was used as the host for cloning work. *A. baylyi* ADP1 (DSM 24193, DSMZ, Germany) was used for all the experiments. The details of all the strains used in the study can be found in Supplemental materials S2.

Modified lysogeny broth (LB) media (10 g/L tryptone, 5 g/L yeast extract, and 1 g/L NaCl) supplemented with 0.4% (w/v) glucose were used to grow *E. coli* and ADP1 for cloning and strain construction. Modified LB media were also used in the experiments for characterization of CRISPRi and the cyclohexanone-controlled induction system, and cell size manipulation by *ftsZ* repression and piperacillin treatment; phosphate buffer (3.88 g/L K_2_HPO_4_ and 1.63 g/L NaH_2_PO_4_) and 0.4% (w/v) glucose were added in the media when double reporter-monomeric superfolder GFP (msfGFP) and mScarlet-strains were used. Modified minimal media MA/9 were used to study *aceA* repression, and mineral salts media (MSM) were used in the experiments for WE production; the carbon sources, including casamino acids, acetate, glucose, and oleic acid, were added as appropriate. The composition of modified MA/9 is 4.40 g/L Na_2_HPO_4_, 3.40 g/L KH_2_PO_4_, 1.00 g/L NH_4_Cl, 0.008 g/L nitrilotriacetic acid, 1.00 g/L NaCl, 240.70 mg/L MgSO_4_, 11.10 mg/L CaCl_2_, 0.50 mg/L FeCl_3_. The composition of MSM is 3.88 g/L K_2_HPO_4_, 1.63 g/L NaH_2_PO_4_, 2.00 g/L (NH_4_)_2_SO_4_, 0.1 g/L MgCl_2_·6H_2_O, 10 mg/L Ethylenediaminetetraacetic acid (EDTA), 2 mg/L ZnSO_4_·7H_2_O, 1 mg/L CaCl_2_·2H_2_O, 5 mg/L FeSO_4_·7H_2_O, 0.2 mg/L Na_2_MoO_4_·2H_2_O, 0.2 mg/L CuSO_4_·5H_2_O, 0.4 mg/L CoCl_2_·6H_2_O, and 1 mg/L MnCl_2_·2H_2_O. Antibiotics were added when needed (15 μg/mL gentamycin, 25 μg/mL chloramphenicol, 50 μg/mL kanamycin, and 50 μg/mL spectinomycin).

### 5.2 Genetic engineering

Molecular cloning was performed using established methods. Transformation and the homologous recombination-based genome editing of ADP1 were carried out as described previously (Santala, Efimova, Kivinen, *et al*., 2011). All the plasmids constructed, primers used in the study, and the corresponding description can be found in Supplemental materials S3 and S4. The sgRNAs used in the study are also included in Supplemental materials S4.

The integration vector pJL1 for replacing ADP1’s prophage site with *dCas9* was constructed by Gibson Assembly. For the construction, the plasmids pdCas9-bacteria (a gift from Stanley Qi, Addgene #44249) (Qi *et al*., 2013), pBAV1C-chn (Luo *et al*., 2019), and pIM1463 (a kind gift from Murin, Addgene #30505) (Murin *et al*., 2012) were used as templates for the amplification of *dCas9, chnR* and the cognate promoter *P*_*chnB*_, and the backbone containing the sequences flanking the prophage site, respectively. The plasmid pJL2 was derived from pJL1 by replacing the RBS for *dCas9*, BBa_0034, with BBa_J61138 using USER cloning. The integration vector pJL3 for the mScarlet gene integration at the prophage site was derived from pJL1 by replacing *dCas9* with the mScarlet gene by USER cloning. The plasmid pJL4 and pJL5 were derived from pJL3 by replacing the RBS BBa_0034 with BBa_J61138 and BBa_J61105, respectively, using USER cloning. The integration vector pJL6 for integration of *msfGFP* at the *poxB* (*ACIAD3381*) site was constructed by amplifying *msfGFP* from the plasmid pBG42 (Zobel *et al*., 2015) and cloning it to the vector pAK400c using restriction sites MunI and XhoI. The plasmid contains sequences flanking *poxB* (Santala, Efimova, Karp, *et al*., 2011) to facilitate genomic integration. The plasmid pJL7 differs from pJL6 in the way that it contains a kanamycin marker instead of a chloramphenicol marker and the sequences flanking *ACIAD3309* instead of *poxB*. The integration vector pJL9 for *acr1* (*ACIAD3383*) knock-out was constructed by cloning a chloramphenicol marker to the pUC57 plasmid containing the sequences flanking *acr1* (here named as pJL8, purchased from Genscript). The plasmid pJL10 was constructed by cloning the *P*_*T5*_-*acr1*-kan^r^ cassette (Luo *et al*., 2020) to pJL8. The plasmid pJL11 contains a null sgRNA under the control of the arabinose inducible promoter, constructed by USER cloning: the plasmids pBAV1C-ara-luxCDE (Santala, Efimova, *et al*., 2014) was used to amplify the backbone as well as *araC* and the cognate promoter *P*_*araBAD*_; the plasmids pSEVA238-CRISPR and pSEVA648 were used to amplify the sgRNA scaffold and the gentamycin resistance gene respectively. Other sgRNA expressing plasmids were derived from pJL11 by USER cloning. The specificity of sgRNAs was checked using Cas-OFFinder (Bae *et al*., 2014). The cassette containing the two-sgRNA array (targeting *msfGFP* and *mScarlet*) and the codon-optimized *cas6* were purchased from Genscript and further cloned to the sgRNA expressing backbone, resulting in pJL17. The plasmid pJL18 was derived from pJL17 by removing *cas6*.

The msfGFP reporter strain (ASA514, also contains the bacterial luciferase operon *luxCDABE* from *Photorhadus luminescens*) was constructed as described below: the strain ADP1 Δ*acr1*::kan^r^/*tdk* (a kind gift from Veronique de Berardinis, Genoscope, France) was transformed with the plasmid pVKK81-T-lux described in (Santala, Karp, *et al*., 2014) for genome integration of *luxCDABE*; the resulting strain was then transformed with pJL6. The msfGFP and mScarlet double reporter strain (ASA518) was constructed by transforming a *mScarlet* expressing strain (Losoi *et al*., 2019) with pJL7. Knock-out of the rod shape-determining genes, *mreB*, and *pbpA*, was done by transformation with the linear cassettes constructed by assembling the corresponding left flanking sequence, a chloramphenicol marker, and the right flanking sequence using SOE-PCR.

### 5.3 Characterization of the cyclohexanone-controlled induction system

The strain ASA511 was used to characterize the cyclohexanone-inducible promoter system, which contains *mScarlet* (integrated into the genome) under the control of the cyclohexanone-inducible promoter. The strain was cultivated in modified LB media (5-mL/14-mL-cultivation tube) containing different concentrations of cyclohexanone at 30 L and 300 rpm. After 25 h, 200 μL of each sample was transferred to a 96-well plate (flat bottom, μClear™, white, Greiner) to measure mScarlet fluorescence and optical density at 600 nm (OD_600_). The measurement was conducted in Spark multimode microplate reader (Tecan, Switzerland); the excitation and emission wavelength were set to 580 and 610 nm. The data of the induction experiment was fit to a model (see Supplemental Note) using the lsqcurvefit function in MATLAB.

### 5.4 Fluorescence measurements for CRISPRi characterization

For the GPF reporter strains, cells were cultivated in 200 μL modified LB media in 96-well plates. The well-plate was incubated in Spark multimode microplate reader at 30 L; double orbital shaking was performed for 5 min twice an hour with an amplitude of 6 mm and a frequency of 54 rpm; OD600 and msfGFP fluorescence signal were measured every 30 min. The excitation and emission wavelength were set to 485 and 535 nm for msfGFP fluorescence measurement. For the msfGFP and mScarlet double reporter strains, cells were cultivated in 4 mL modified LB media in 14-mL-cultivation tubes, and phosphate buffer and 0.4% (w/v) glucose were also added to the media. The cultivation was conducted in a shaker at 30 L and 300 rpm. Samples were taken at 20 h for fluorescence measurement. The expressions of *dCas9* and sgRNAs were induced at the beginning of the cultivation by adding 5 μM cyclohexanone and 1% (w/v) arabinose, respectively.

### 5.5 Cultivations for repression studies and WE production

To study the repression of *aceA* by CRISPRi, cells were cultivated in 20 ml modified MA/9 media in 100-mL-flasks at 30 L and 300 rpm for 20 h; the media were supplemented with 0.2% (w/v) casamino acids and 40 mM acetate. Inducers were added at the beginning of the cultivation. For the *ftsZ* repression study, cells were cultivated in 5 mL modified LB media in 14-mL-cultivation tubes at 30 L and 300 rpm. Inducers were added when OD600 reached 0.2, after which the cultivation continued for 18 h. Induction was performed by adding 5 μM cyclohexanone and 1% (w/v) arabinose.

For WE production, cells were cultivated in 250-mL-flasks containing 50 ml MSM supplemented with 0.2% (w/v) casamino acids and different carbon sources. The cultivation was performed in three different conditions. In this first condition, 5 mM glucose and 60 mM acetate were added at the beginning and supplied twice at ∼10 h and ∼21 h with a concentration of 5 mM and 60 mM, respectively. The cultivation was conducted at 30 L and 300 rpm with an initial OD of 0.1. Inducers were added into parallel cultures when OD reached ∼0.5 (early induction) and ∼1.5 (late induction), respectively. In the second condition, 200 mM glucose was added, and cells were cultivated at 25 L and 300 rpm. The initial OD was 0.1, and inducers were added into parallel cultures when OD reached ∼1.5 (early induction) and ∼4 (late induction), respectively. In the third condition, 10 mM glucose and 14 g/L oleic acid (65.0-88.0%, Sigma Aldrich) were supplemented. Cells were cultivated at 30 L and 300 rpm with an initial OD of 0.05. After 12 h, 10 mM glucose was supplied, and inducers were added. Induction was performed with 1% (w/v) arabinose and 50 μM cyclohexanone.

### 5.6 Analysis of cell size by microscopy

Cells were harvested by centrifugation at 8000 rpm for 10 min and resuspended and diluted with phosphate-buffered saline (PBS). The cell ODs after dilution ranged from 0.05 to 0.2. The cells (5-10 μL) were then heat-fixed on a microscope slide and stained with crystal violet staining reagent for one minute and subsequently washed with tap water for several seconds. The cells were analyzed under bright field illumination using Zeiss Axioskop 2 equipped with Achroplan 100x objective lens. Image processing and cell size analysis were done using the software Fiji-ImageJ (Schindelin *et al*., 2012). Briefly, images were processed through the following steps: 8-bit conversion, background subtraction, thresholding, hole filling (if needed), and particle analysis; any defect was manually removed. For each sample, 200-300 cells were taken randomly for cell size analysis based on the cross-section areas using RStudio.

### 5.7 Cell sedimentation analysis

Cells were harvested by centrifugation and resuspended with PBS in glass test tubes. All samples were diluted to an OD600 of 0.9 and left undisturbed at room temperature. The OD600 was measured periodically using the spectrophotometer Ultrospec 500pro (GE Healthcare Life Sciences, USA).

### 5.8 Nile red staining

Nile red staining was applied to visualize neutral lipid bodies (Wältermann *et al*., 2005). Specifically, Nile red stock solution (0.5 mg/mL DMSO) was diluted to 0.5 μg/mL using PBS; cells were harvested by centrifugation at 12000 rpm for 5 min; the cells were then resuspended with the diluted Nile red solution and incubated on ice for 30 min; after staining, cells were washed with PBS and used for visualization under the microscope with an appropriate filter.

### 5.9 Analysis of substrate consumption

Cell cultures were centrifuged at 12000 rpm for 5 min, and the supernatant was collected and filtered with 0.2 μm pore size filters. The samples were further diluted with water and subjected to analysis with high-performance liquid chromatography (HPLC). The analysis was conducted with an LC-20AC prominence liquid chromatograph (Shimadzu, USA) equipped with RID-10A refractive Index detector. Phenomenex Rezex RHM-monosaccharide H+ (8%) column (Phenomenex, USA) was used, and 5 mM sulfuric acid was used as a mobile phase with a flow of 0.4 mL/min.

### 5.10 Analysis of WE production

WEs were visualized using thin-layer chromatography (TLC), as described before (Santala, Efimova, Kivinen, *et al*., 2011). Briefly, 1-3 mL of biomass was harvested from cultures by centrifugation. The cell pellets were resuspended in 500 μL methanol and vortexed for 30 min. Then, 250 μL chloroform was added, after which the samples were vortexed for 1 h. Then the samples were centrifuged again. Next, 250 μL chloroform was 250 μL PBS were added, and the samples were slowly swirled for 2 h. After that, centrifugation was applied, and the low phase was taken for TLC. Glass HPTLC Silica Gel 60 F_254_ plate (Merck, USA) was used for TLC. The mobile phase was a mixture of hexane, diethyl ether, and acetic acid with a ratio of 90:15:1. Jojoba oil was used as the standard of WEs. The TLC plats were stained with iodine.

Quantification of WEs was carried out using ^1^H nuclear magnetic resonance (NMR). The cells were harvested from 40 mL culture by centrifugation at 30000 *g* for 30 min. The harvested cells were freeze-dried with Christ APLHA1-4 LD plus freeze dryer (Germany) for 24 h. The extraction of lipids from the freeze-dried cells and analysis with NMR were performed as described previously (Santala, Efimova, Karp, *et al*., 2011). The chemical shift of 4.05 ppm corresponds to the proton signal of α-alkoxy-methylene group of WEs. The integral value of the signal at 4.05 ppm was used to calculate the content of WEs. An average molar mass of 506 g/mol was used to calculate the WE titer in grams per liter (Lehtinen *et al*., 2018).

## Supporting information

Supplemental materials S2 (strains)

Supplemental materials S3 (plasmids)

Supplemental materials S4 (primers and sgRNAs)

## Acknowledgements

Funding: The work presented in this article is supported by Academy of Finland (grants no. 334822, 310188) and The Novo Nordisk Foundation grant NNF21OC0067758. In addition, the financial support from The Novo Nordisk Foundation through grants NNF20CC0035580 and TARGET (NNF21OC0067996), and the European Union’s Horizon 2020 Research and Innovation Programme under grant agreement No. 814418 (SinFonia) to the Systems Environmental Microbiology laboratory is gratefully acknowledged. Part of this work was carried out by S. Santala in the Systems Environmental Microbiology laboratory at The Novo Nordisk Foundation for Biosustainability in the context of a research stay supported by Academy of Finland (grant no. 310135).

## Author contribution

Author contribution: JL, SS, and VS designed the study. JL, SS, EE, and DCV carried out the research work. JL, SS, VS, and DCV analyzed the data. SS and VS supervised the study. All authors participated in writing the manuscript. All authors read and approved the final manuscript.

