## Supplemental materials S1 for "Engineering cell morphology by CRISPR interference in *Acinetobacter baylyi* ADP1"

Jin Luo^1^*, Elena Efimova^1^, Daniel Christoph Volke^2^, Ville Santala^1^, Suvi Santala^1^
^1^Faculty of Engineering and Natural Sciences, Hervanta campus, Tampere University, Korkeakoulunkatu 8, Tampere, 33720, Finland

^2^ The Novo Nordisk Foundation Center for Biosustainability, Technical University of Denmark, Kemitorvet 220, 2800 Kgs. Lyngby, Denmark

*Corresponding author

Email addresses:

Jin Luo:
Elena Efimova:
Daniel Christoph Volke:
Ville Santala:
Suvi Santala:

**Supplemental Materials**

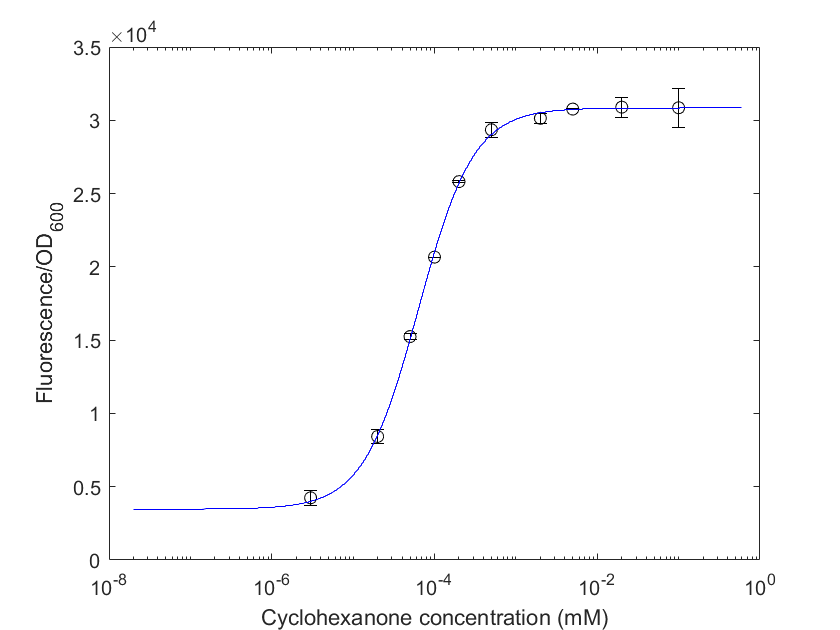

**Figure S1.** Transfer function of the cyclohexane-inducible promoter. The transfer function shows how the activity of the cyclohexane-inducible promoter (expressed as mScarlet fluorescence/OD_600_) changes with cyclohexane concentration. Data represent average values ± standard deviations of two independent biological experiments. The blue line is the fit to the data points using the model shown in equation 3 (see Supplemental Note).

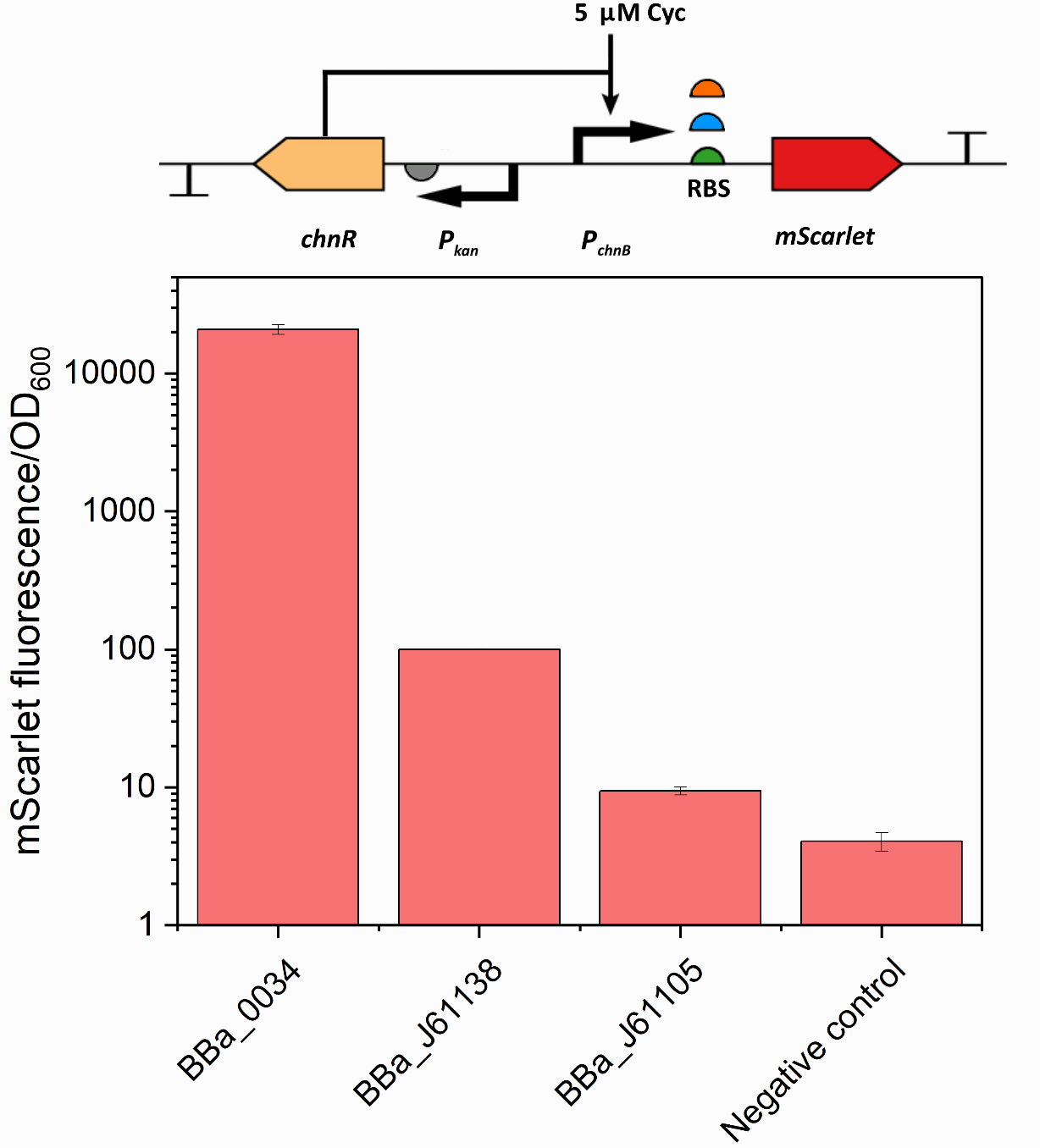

**Figure S2.** Comparison between the strength of three ribosome binding sites (RBSs) in ADP1. The top panel shows the architecture of the reporter construct. The three RBSs, BBa_0034, BBa_J61138, and BBa_J61105, are upstream of *mScarlet*, which is under the control of the cyclohexanone-inducible promoter. The cassette was integrated into the genome. Strains were grown in LB media containing 5 µM cyclohexanone (Cyc). Samples were taken for fluorescence measurement after 24 h. Data represent average values ± standard deviations of two independent biological experiments.

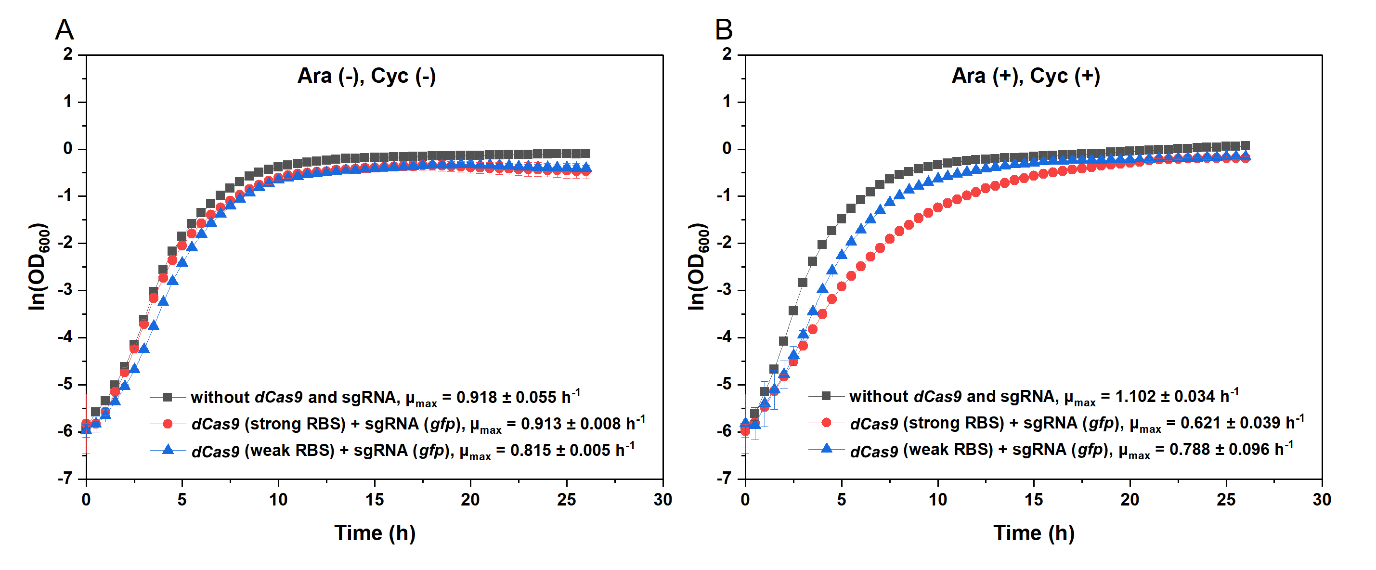

**Figure S3.** The effect of *dCas9* expression level on growth rate. The growth in non-inducing (A) and inducing (B) conditions are shown. The two strains ASA515 and ASA516 contain different RBSs preceding the *dCas9* gene: either the strong BBa_0034 or the weak BBa_J61138. The sgRNA targets the constitutively expressed *gfp*. The reporter strain ASA514 without the CRISPRi machinery was used as a control. Cells were cultivated in LB media at 30 ℃. For induction, 1% (w/v) arabinose (Ara) and 5 µM cyclohexanone (Cyc) were added in the beginning. The maximal specific growth rate (µ_max_) was calculated by taking the slope for the linear part of the logarithmic OD_600_ (R^2^ >0.99). Data represent average values ± standard deviations of two independent biological experiments.

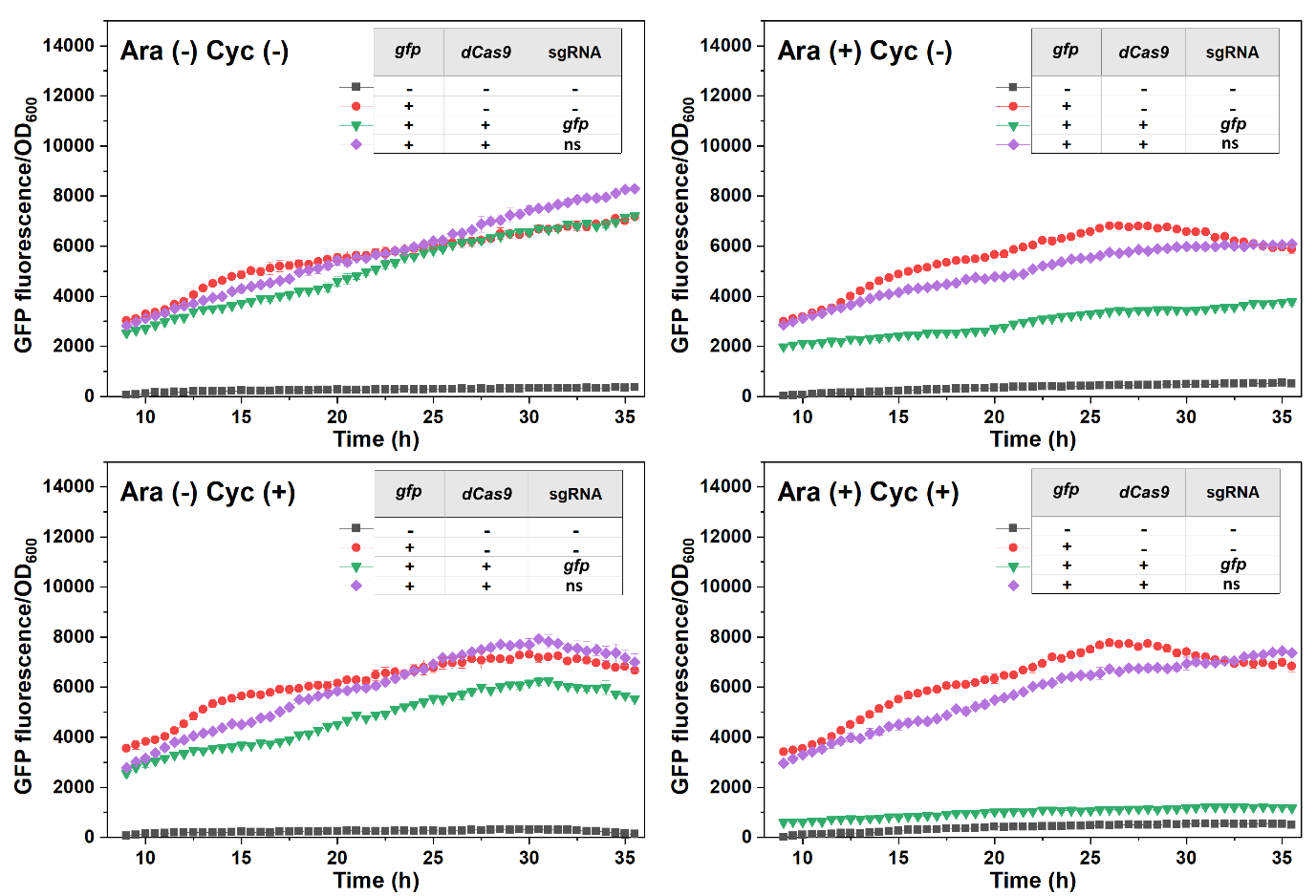

**Figure S4.** Change of GFP fluorescence/OD_600_ during cultivation for each ADP1 strain under different induction states. Cells were grown in LB media and cultivated at 30 ℃. For induction, 1% (w/v) arabinose (Ara) and/or 5 µM cyclohexanone (Cyc) were supplemented. The non-specific (ns) sgRNA was used as a control. Data represent average values ± standard deviations of two independent biological experiments.

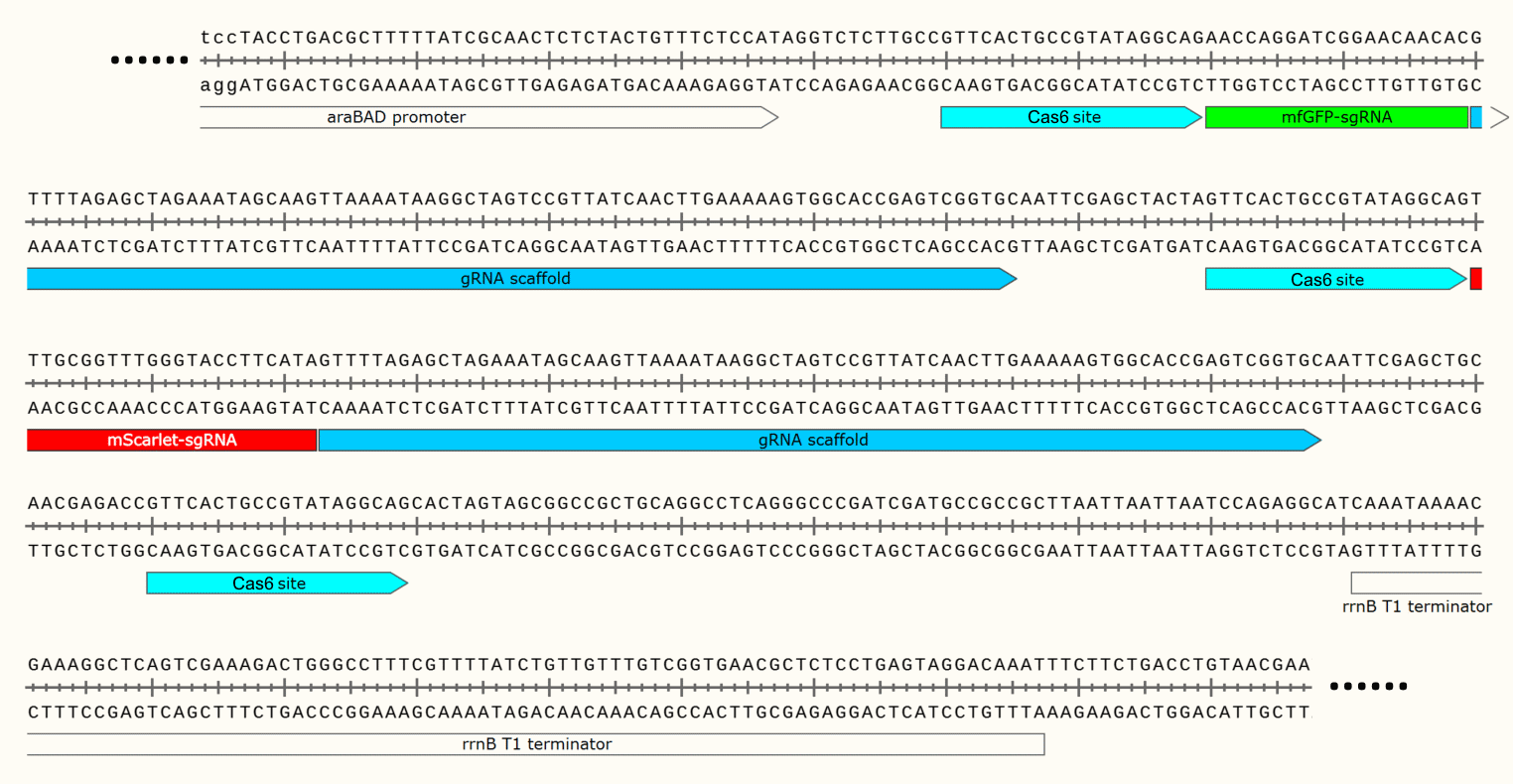

**Figure S5.** DNA sequence of the guide RNA array containing the *gfp*-targeting sgRNA and the *mScarlet*-targeting sgRNA.

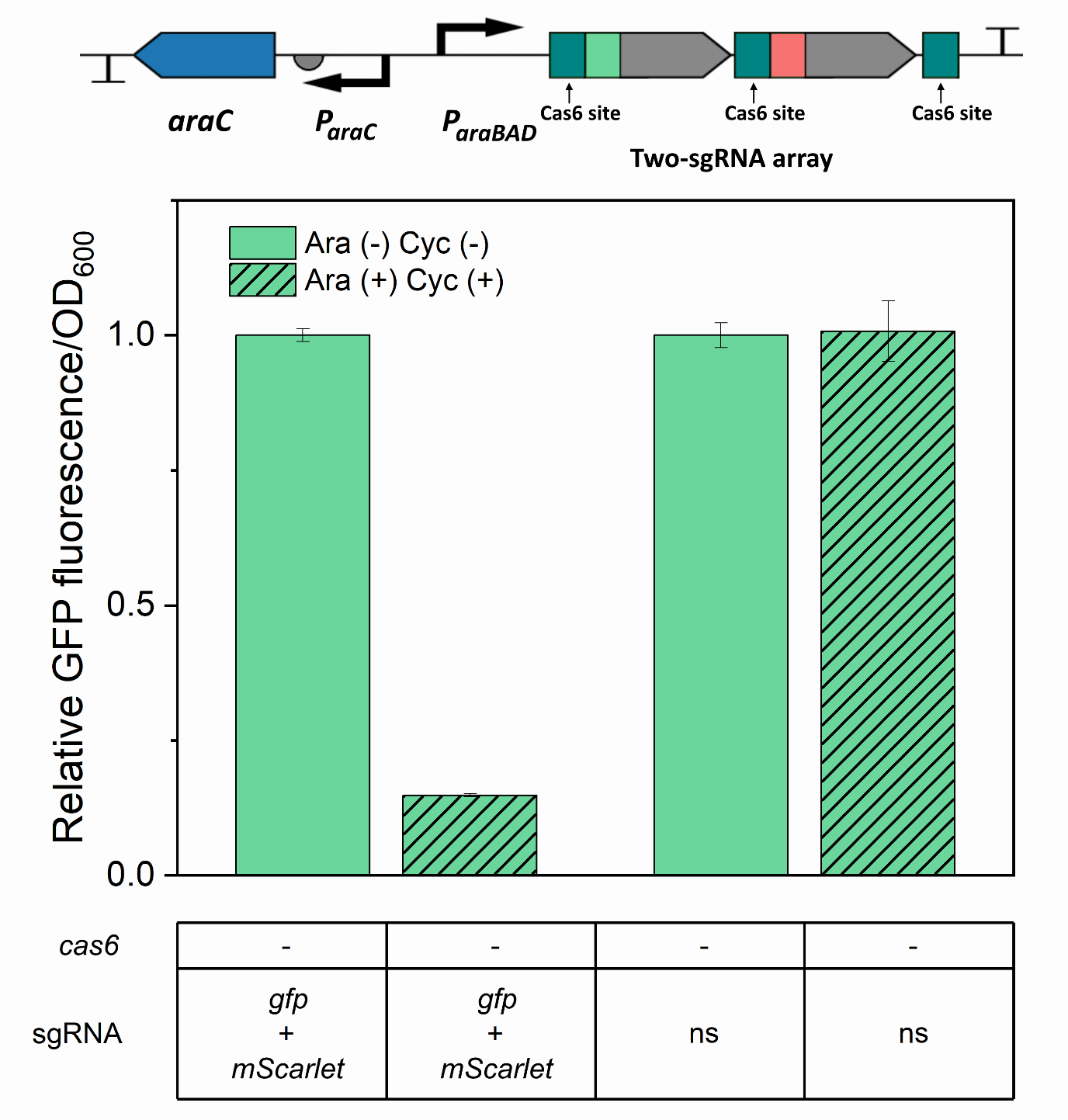

**Figure S6.** Validation of GFP repression by expression of the two-sgRNA array without *cas6*. To further validate that *msfGFP*-targeting sgRNA, which was placed before the *mScarlet*-targeting sgRNA in the two-sgRNA array (as shown in the top panel), was still functional without *cas6*, the cassette (integrated into the pBAV plasmid) was introduced into the GFP single reporter strain containing *dCas9*. The GFP reporter strain containing *dCas9* and a non-specific (ns) sgRNA was used as control. Cells were grown in buffered LB containing 0.4% (w/v) glucose and cultivated at 30 ℃. For induction, 1% (w/v) arabinose (Ara) and 5 µM cyclohexanone (Cyc) were supplemented. Samples were taken for fluorescence measurement at 20 h. The fluorescence values for each strain were normalized to the values measured for this strain without induction. Data represent average values ± standard deviations of two independent biological experiments.

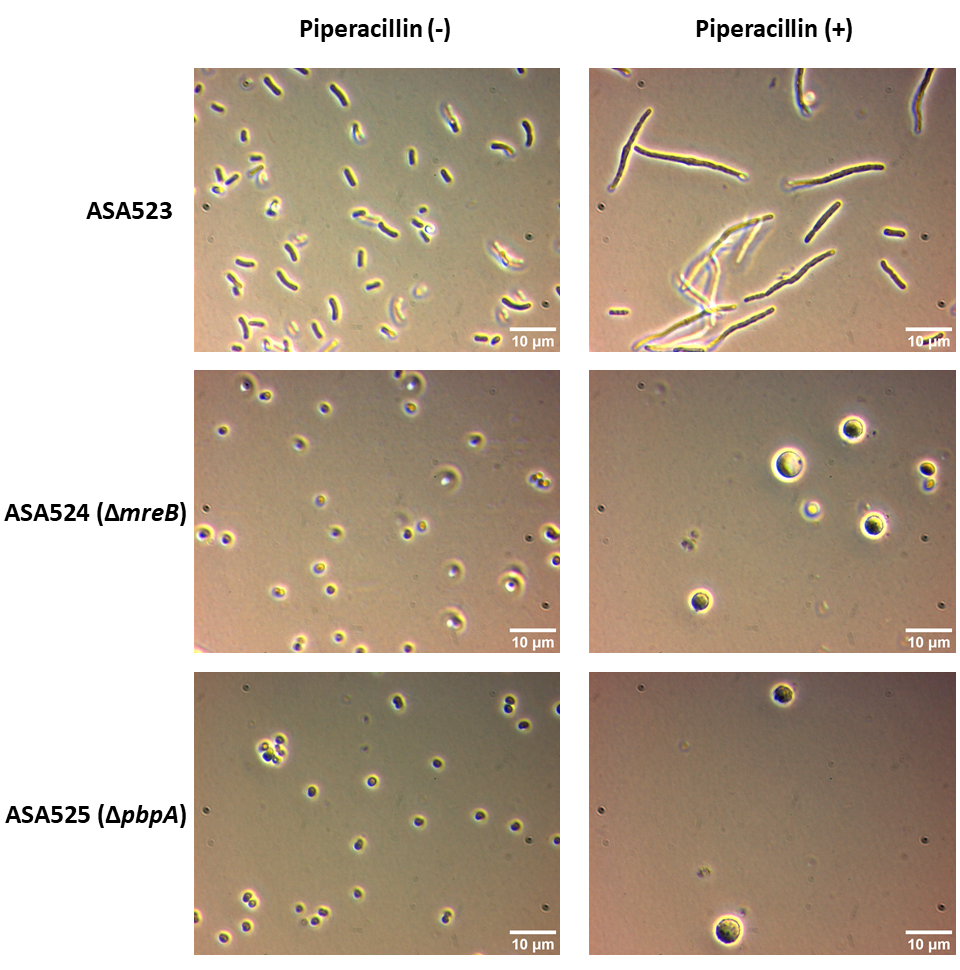

**Figure S7.** Cells treated with piperacillin were enlarged. Cells were cultivated in MA/9 media supplemented with 200 mM glucose at 25 ℃ for 20 h. Piperacillin with a final concentration of 12 µg/mL was used for the treatment. The strain ASA523 contained a copy of acr1 under the control of the constitutive promoter T5. ASA524 and ASA525 were derived from ASA523 by deleting the rod shape-related genes mreB and pbpA respectively.

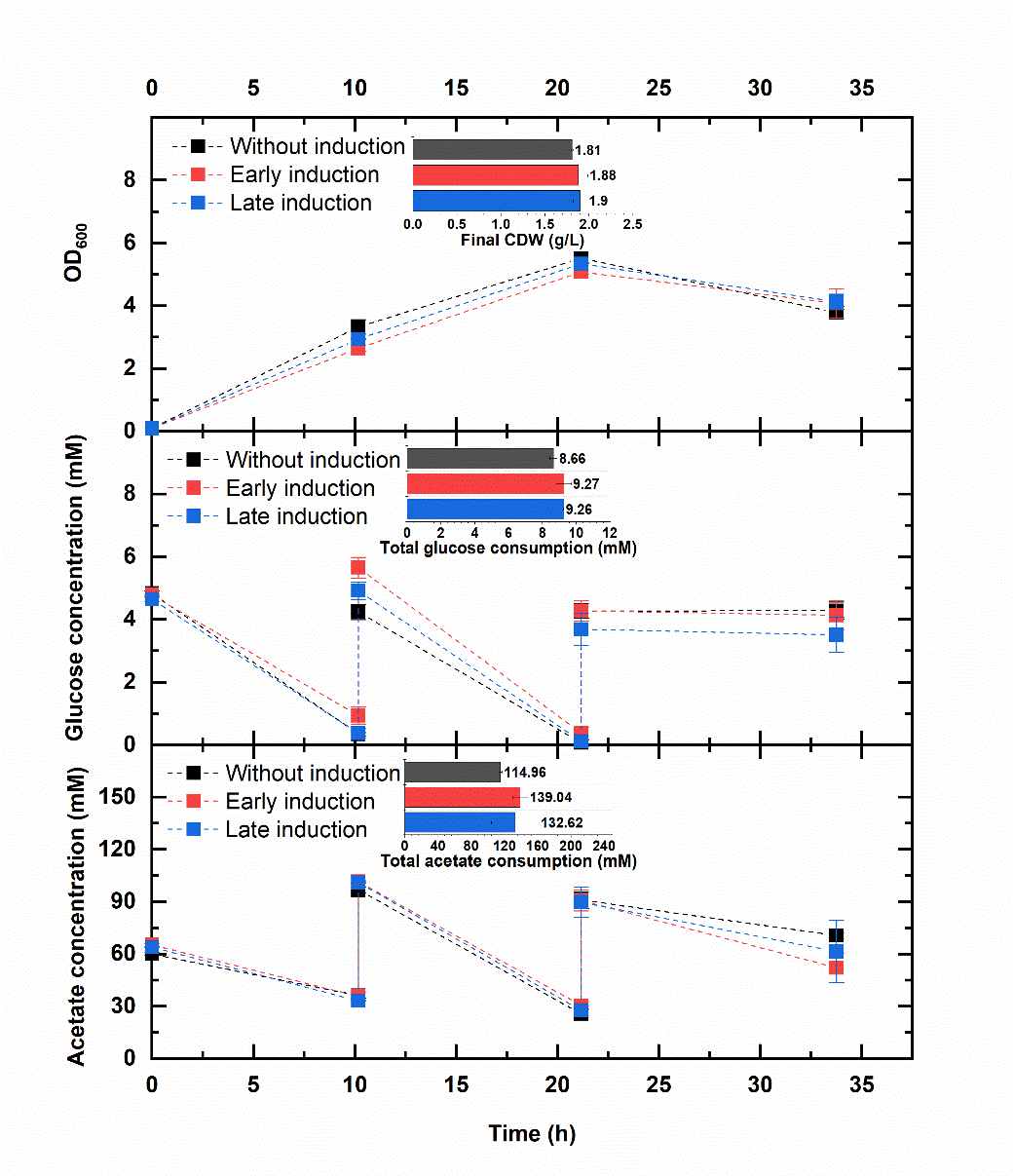

**Figure S8.** Cultivation of ASA531 (*aceA* deletion + *acr1* overexpression + CRISPRi targeting *ftsZ*) in glucose and acetate. Cells were grown in mineral salts media containing 0.2% casamino acids, 5 mM glucose, and 60 mM acetate, and cultivated at 30 ℃. Glucose (5 mM ) and acetate (60 mM) were fed again at 10 h and 21 h. Two parallel cultivations were performed, and inducers (1% (w/v) arabinose and 50 µM cyclohexanone) were added when the OD reached ~0.5 (early induction) and ~1.5 (late induction) respectively. The OD (top panel), glucose concentration (middle panel), and acetate concentration (bottom panel) were measured at 0, 10, 21, and 34 h. Data represent average values ± standard deviations of two independent biological experiments.

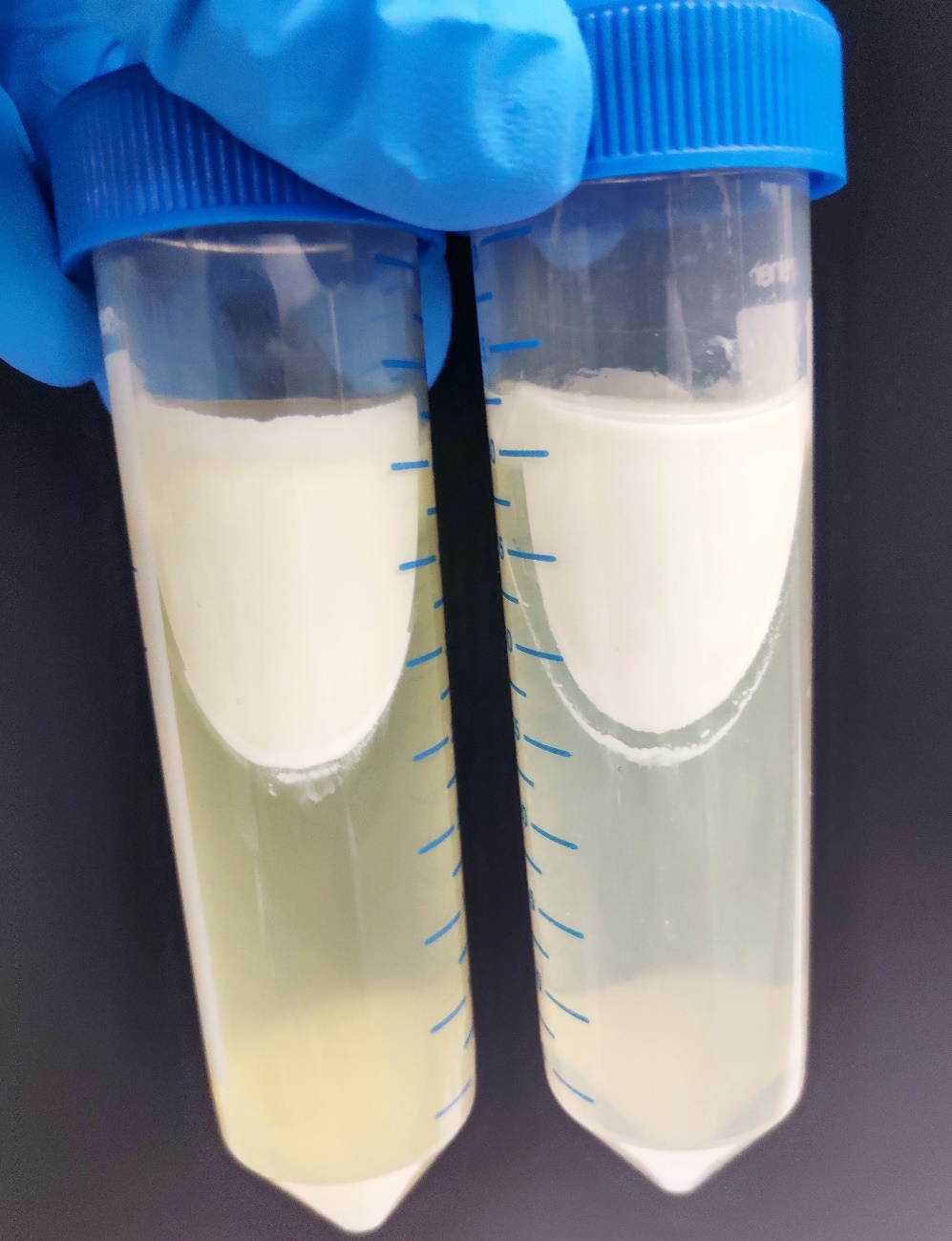

**Figure S9.** Cultures after centrifugation for harvesting. Part of the culture was floating or suspended in the media after centrifugation (12000 rpm for 2.5 h) when fatty acids were used as the substrate for the cultivation.

**Supplemental Note**

**Mathematical analysis for the cyclohexanone induction system**

The regulator ChnR was reported to be an activator (Iwaki et al., 1999; Steigedal and Valla, 2008), in response to cyclohexanone. The binding of cyclohexanone (Cyc) to ChnR is described with equation 1,

$$\text{F}\text{ChnR:Cyc}=\frac{\left[ \mathrm{Cyc} \right]^{n}}{K_{d}^{n}+\left[ \mathrm{Cyc} \right]^{n}} \left( 1 \right)$$

where F_chnR:Cyc_ is the fraction of ChnR bound to cyclohexanone, [Cyc] is cyclohexanone concentration, K_d_ is the dissociation constant, and n is the cooperativity. The activity of the promoter is described with equation 2,

$$\text{P}\text{chnB}=P_{\mathrm{chnB}}^{\max}*\frac{\text{K}\text{1}\text{+K}\text{2}*\text{F}\text{ChnR:Cyc}}{1+\text{K}\text{1}\text{+K}\text{2}*\text{F}\text{ChnR:Cyc}} \left( 2 \right)$$

where P_chnB_ is the activity of the cyclohexanone-inducible promoter, Pis the maximal promoter activity. K_1_and K_2_ are the terms of two states of the promoter (Figure S11). As the promoter activity can be reflected by the fluorescence signal, the fluorescence signal can be described using a similar equation below,

$$\left[ fluorescence/OD \right]=\left[ fluorescence/OD \right]_{\max}*\frac{\text{K}\text{1}\text{+K}\text{2}*\text{F}\text{ChnR:Cyc}}{1+\text{K}\text{1}\text{+K}\text{2}*\text{F}\text{ChnR:Cyc}} (3)$$

Where [fluorescence/OD] is the measured fluorescence per OD, [fluorescence/OD]_max_ is the maximal fluorescence signal. The parameters for fitting the model to the experimental data are shown in Table S1.

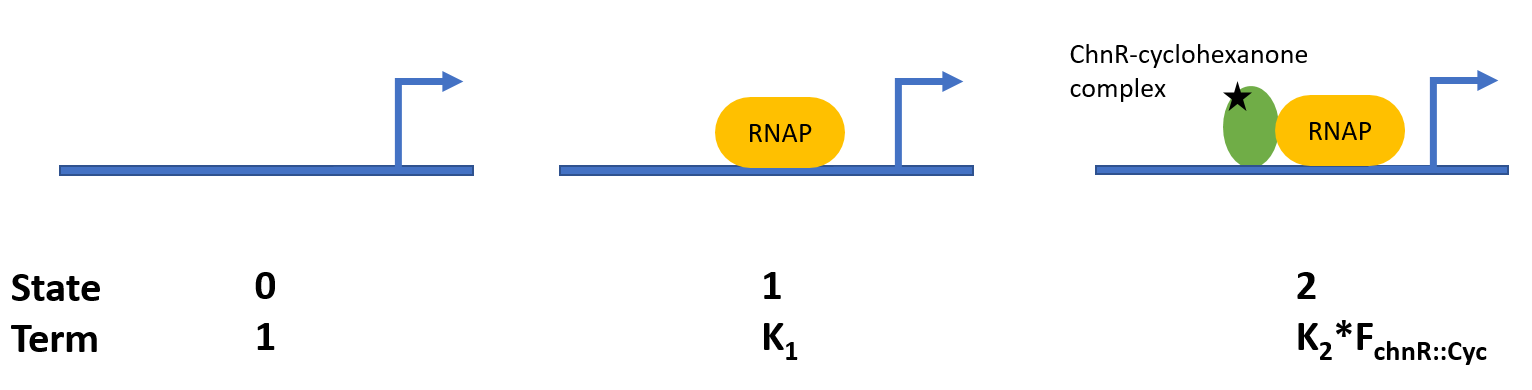
**Figure S11.** A model assuming three major states of the cyclohexanone-inducible promoter. In the first state, the promoter is not occupied. In the second state, the promoter binds to RNA polymerase. In the third state, the promoter is activated, binding to the ChnR-cyclohexanone complex and RNA polymerase. The amounts of RNA polymerase and ChnR are assumed to be constant.

**Table S1.** Parameters for data fitting.

| [fluorescence/OD]_max_ | K_1_ | K_2_ | K_d_ | n |
| --- | --- | --- | --- | --- |
| 31718 | 0.12 | 35.02 | 0.00095 | 1.28 |
